# Broadly conserved FlgV controls flagellar assembly and *Borrelia burgdorferi* dissemination in mice

**DOI:** 10.1101/2024.01.09.574855

**Authors:** Maxime Zamba-Campero, Daniel Soliman, Huaxin Yu, Amanda G. Lasseter, Yuen-Yan Chang, Jun Liu, L. Aravind, Mollie W. Jewett, Gisela Storz, Philip P. Adams

## Abstract

Flagella propel pathogens through their environments yet are expensive to synthesize and are immunogenic. Thus, complex hierarchical regulatory networks control flagellar gene expression. Spirochetes are highly motile bacteria, but peculiarly in the Lyme spirochete *Borrelia burgdorferi*, the archetypal flagellar regulator σ^28^ is absent. We rediscovered gene *bb0268* in *B. burgdorferi* as *flgV,* a broadly-conserved gene in the flagellar superoperon alongside σ^28^ in many Spirochaetes, Firmicutes and other phyla, with distant homologs in Epsilonproteobacteria. We found that *B. burgdorferi* FlgV is localized within flagellar motors. *B. burgdorferi* lacking *flgV* construct fewer and shorter flagellar filaments and are defective in cell division and motility. During the enzootic cycle, *B. burgdorferi* lacking *flgV* survive and replicate in *Ixodes* ticks but are attenuated for dissemination and infection in mice. Our work defines infection timepoints when spirochete motility is most crucial and implicates FlgV as a broadly distributed structural flagellar component that modulates flagellar assembly.

## INTRODUCTION

Flagella are complex membrane-embedded nanomachines with a protruding filament that allow bacteria to move through liquids and across solid surfaces. Across microorganisms, flagella are critical for organisms to inhabit different environmental niches, to migrate toward nutrient-rich regions, to escape unfavorable conditions, and to move from host to host. For several bacteria, flagella are even connected to basic processes such as cell division and maintenance of cell shape (reviewed in (Wolgemuth *et al*., 2006, Yang *et al*., 2016, Popp & Erhardt, 2021). Bacteria in the phylum Spirochaetota are distinguished by unique endoflagella, which are anchored at each cell pole and extend through the periplasmic space, rather than the extracellular environment (reviewed in (Wolgemuth, 2015, San Martin *et al*., 2023)). It is this flagellar placement that creates the flat-wave structure of the spirochetal bacterium *Borrelia* (*Borreliella*) *burgdorferi* (Motaleb *et al*., 2000), the Lyme disease pathogen.

Lyme disease is an emerging infectious disease and the foremost vector-borne illness in the United States, with almost half a million infections estimated annually (Kugeler *et al*., 2021). *B. burgdorferi* exists in a complex enzootic cycle that requires acquisition and transmission of the spirochete between *Ixodes scapularis* ticks and small mammals and birds (reviewed in (Radolf *et al*., 2012)). Motility of *B. burgdorferi* is essential for survival throughout the enzootic cycle (Sultan *et al*., 2013). Larval ticks are hatched uninfected and acquire *B. burgdorferi* upon feeding on an infected host, most commonly the white-footed mouse in the northeastern United States. During larval-tick feeding, the bacterium navigates from the mouse dermis to colonize the tick midgut. The tick will then molt into a nymph, infected with *B. burgdorferi*. Inside the unfed nymph, spirochetes remain sessile to survive the nutrient-poor midgut and await the next bloodmeal, which can occur months to a year after the molt. When the nymphal tick bites its next host, spirochetes associate with the basement membrane of the midgut as nonmotile aggregates and then become motile, migrating from the tick midgut to the tick salivary glands (Dunham-Ems *et al*., 2009), ultimately infecting the skin of a mammalian host. In the mammal, *B. burgdorferi* spread from the initial bite site through the blood (hematogenous dissemination) and other routes, to colonize distal tissues. Overall, to progress through this enzootic cycle, *B. burgdorferi* must finely tune its motility to conserve energy when needed, disseminate when able, and overcome host immune responses.

Regulation of bacterial motility, which can be at the level of gene expression of flagellar proteins or the functional level of flagellar motor rotation and chemotaxis, has been studied most in Enterobacteriaceae and Bacillaceae, (reviewed in (Soutourina & Bertin, 2003, Chevance & Hughes, 2008, Guttenplan & Kearns, 2013)). In these model organisms, the genes encoding the basal body, rod and hook of the nascent flagellum and the alternative sigma factor, σ^28^ (σ^F^, FliA, SigD), are transcribed first. Once the hook–basal body is assembled, the genes encoding the flagellar filament are transcribed, controlled by σ^28^ associating with RNA polymerase. This hierarchical gene regulation prevents malformed hook–basal body complexes and allows a bacterium to rapidly control motility in an energetically efficient way. The genus *Borrelia/Borelliella*, but not other spirochetes (e.g., *Treponema*, *Spirochaeta*, *Leptospira*, and *Turneriella*) lacks the gene encoding σ^28^. Thus, all *B. burgdorferi* flagellar genes are reported to be transcribed constitutively under the control of the housekeeping sigma factor, σ^70^, signifying the first bacterium discovered to have this scheme of flagellar gene expression (Ge *et al*., 1997a, Ge *et al*., 1997b).

The conserved and generally recognized motility “superoperon” is denoted the “*flgB* operon” in *B. burgdorferi* and includes 31 genes (*bb0294* to *bb0264*) (Ge *et al*., 1997b, Zhang *et al*., 2020). These genes encode the components of the flagellar structure (basal body, hook, rod and filaments), motor ATPases, and flagellar assembly proteins. Cryo-electron tomography has visualized the *in situ* flagellar motor structure and sequential assembly of *B. burgdorferi* periplasmic flagella (Zhao *et al*., 2013), yet the regulation of this temporal assembly and how environmental signals impact flagellar biosynthesis remain poorly understood. Recent work in *B. burgdorferi* identified FlhF (BB0270) as a potential flagellar regulator encoded within the *flgB* operon (Zhang *et al*., 2020). FlhF is a GTPase that influences the number and position of *B. burgdorferi* flagella. This protein is well studied in other bacteria, where it interacts with FlhG (also named FleN and MotR; a MinD-like ATPase) and flagellar transcription factors to facilitate flagellar assembly (reviewed in (Schuhmacher *et al*., 2015)). *B. burgdorferi* encodes an FlhG homolog (BB0269), directly downstream of *flhF* (*bb0270*), but the exact mechanism of *B. burgdorferi* FlhF- and FlhG*-*mediated flagellar regulation has yet to be determined.

In this study we characterize another gene in the *flgB* superoperon, *bb0268*. BB0268 was previously annotated as a homologue of Hfq (Lybecker et al., 2010). Hfq, which is best characterized in Enterobacteriaceae, is a homohexameric RNA binding protein of the Sm domain superfamily that stabilizes small RNAs (sRNAs) and facilitates sRNA base pairing with cognate RNA targets in many bacteria (reviewed in (Updegrove *et al*., 2016, Santiago-Frangos & Woodson, 2018)). In our recharacterization of *bb0268*, we demonstrated that the gene is evolutionarily unrelated to Hfq. BB0268 has an entirely different structure from Hfq and does not bind RNA. Comparative genomics analysis of Hfq suggests *B. burgdorferi* lacks an Hfq homolog entirely. Instead, we found strong co-conservation of *bb0268* homologs with the genes encoding FlhF, FlhG, and σ^28^, spanning the genomes of numerous flagellated bacteria, and thus we renamed *bb0268* as *flgV*. We discovered FlgV localizes within the *B. burgdorferi* flagellar motor and impacts the number and length of flagellar filaments, as well as *B. burgdorferi* dissemination during mouse infection. We therefore propose that FlgV plays previously uncharacterized structural and functional roles in late-stage flagellar assembly.

## RESULTS

### BB0268 was misannotated as an RNA-binding protein

*B. burgdorferi* harbors a large 26.3 kb “*flgB* operon” which encompasses 31 flagellar biosynthesis genes, including *flhF* (*bb0270*) and *flhG* (*bb0269*), potential regulators of flagellar biosynthesis (Zhang *et al*., 2020). One study previously suggested that *bb0268* (present directly downstream of *flhF* and *flhG*) encodes an “atypical Hfq homolog” (Lybecker *et al*., 2010). Hfq, an ortholog of the Sm proteins found in archaea and eukaryotes (Zhang *et al*., 2002), is the most-extensively studied bacterial regulatory RNA-binding protein. In many bacteria, the Hfq hexamer uses distinct surfaces to bind specific sequence motifs in small RNAs (sRNAs) and their target RNAs, often resulting in regulatory consequences (reviewed in (Updegrove *et al*., 2016, Santiago-Frangos & Woodson, 2018)).

Given the report that BB0268 is an Hfq homolog, we sought to identify possible RNAs associated with BB0268. We assayed the ability of BB0268 to bind RNA *in vivo* by performing a co-immunoprecipitation (co-IP) experiment with *B. burgdorferi* harboring a C-terminal 3XFLAG tagged derivative of BB0268, expressed from the endogenous chromosomal location. The parent wild-type (WT) strain was used as a control. No growth defect was observed in the BB0268-3XFLAG strain compared to WT (Figure S1A). Additionally, in the same experiment, we used *E. coli* cells expressing a single-FLAG-tagged version of the *E. coli* Hfq (Melamed *et al*., 2020), also encoded at the endogenous locus, and the corresponding *E. coli* WT strain. The tagged proteins were immunoprecipitated (IP) and RNA was isolated from the IP samples. Immunoblot analysis of cell lysate (total), supernatant, and elution (IP) samples confirmed the production of both BB0268-3XFLAG and *E. coli* Hfq-FLAG, which were enriched by IP (Figure 1A). The RNA isolated from the IP samples was analyzed on an Agilent TapeStation system (Figure 1B). Insignificant RNA levels were detected with BB0268-3XFLAG IP compared to ∼17 ng/μl IP RNA with *E. coli* Hfq-FLAG.

**Figure 1.**
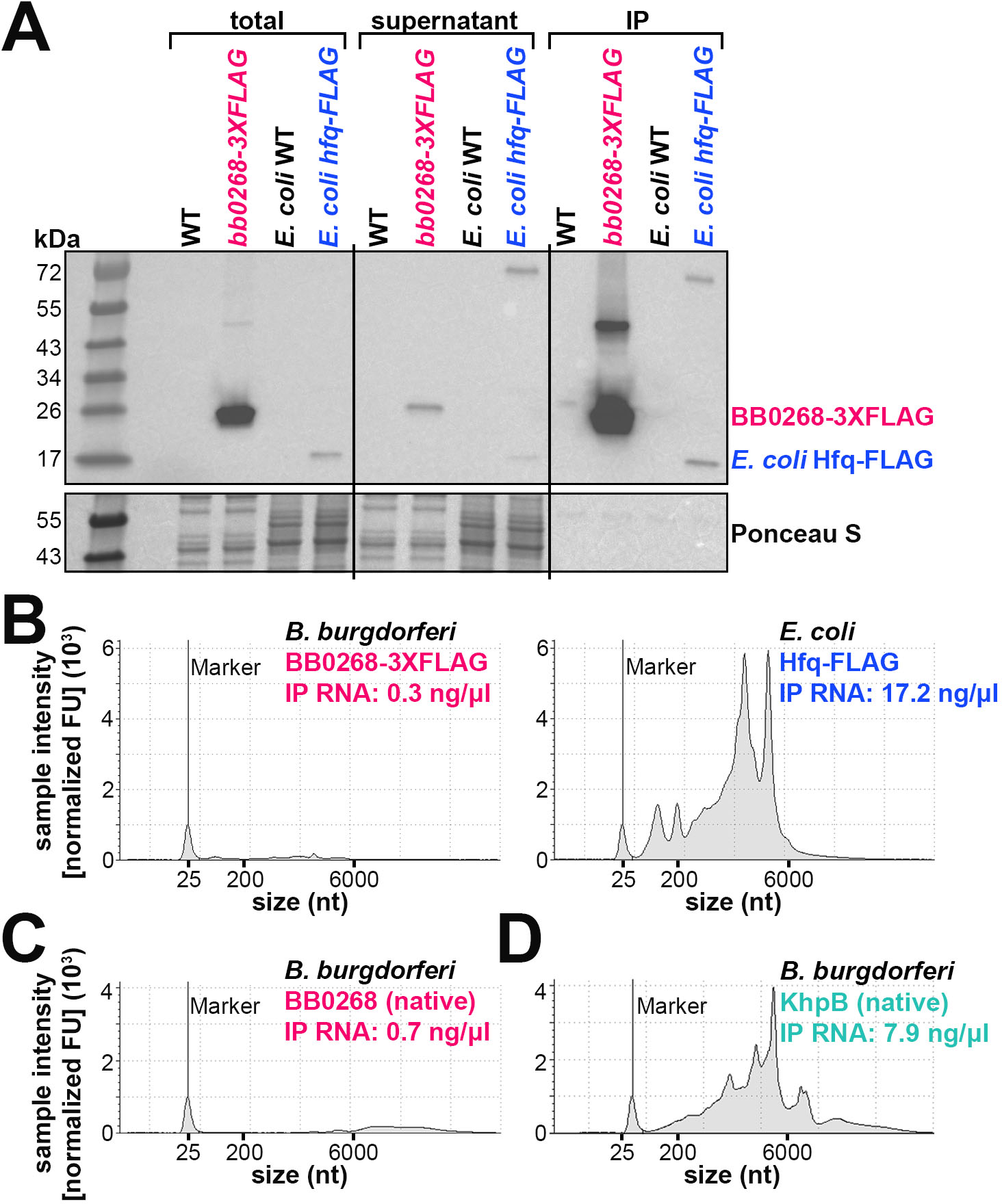
BB0268 does not bind RNA. (**A**) Immunoblot analysis of immunoprecipitated *B. burgdorferi* BB0268-3XFLAG (PA007) or *E. coli* Hfq-FLAG (EC153) samples. The parent WT strains (PA003 and EC152, respectively) were used as controls. *B. burgdorferi* cells were grown to 1×10^8^ cells/ml and *E. coli* cells were grown to an OD_600_ = 1.0 and then exposed to UV to crosslink any RNAs associated with the proteins. After cell lysis, the tagged proteins were immunoprecipitated (IP) and RNA was isolated from the IP samples. Equal volumes of cell lysis (total), supernatant of the α-FLAG-beads during washing, and elution from the α-FLAG-beads (IP) were separated on a Tris-Glycine gel, transferred to a membrane, stained with Ponceau S, and probed using ɑ-FLAG antibodies. Size markers are indicated. **(B)** Quantification of α-FLAG immunoprecipitated RNA. Each sample (2 µl) was analyzed using an Agilent 4200 TapeStation System and the RNA high sensitivity reagents. **(C)** Quantification of α-BB0268 immunoprecipitated RNA, as in panel B. **(D)** Quantification of α-KhpB immunoprecipitated RNA, as in panel B. Panels A and B are a representative experiment from three independent experimental replicates, also including lysates from stationary phase *B. burgdorferi*. All replicates showed the same results.

In a separate experiment, we used BB0268 antibodies (generated for this study; see Materials and Methods) to immunoprecipitate native BB0268. Again, we observed enrichment of BB0268 in the IP fraction by immunoblot analysis (Figure S1B), but no RNA was detected (Figure 1C).

To demonstrate that this co-IP approach can successfully enrich for RNA associated with a *bona fide* RNA-binding protein in *B. burgdorferi* we assayed the KH-domain protein, KhpB. KH domain proteins are another class of RNA-binding proteins in bacteria, and *B. burgdorferi* harbors KhpA (BB0696) and KhpB (BB0443) homologs (reviewed in (Olejniczak *et al*., 2021)). We observed enrichment of KhpB in the IP fraction by immunoblot analysis with antibodies targeting native KhpB (Figure S1C). In contrast to BB0268, we detected ∼8 ng/μl IP RNA with KhpB (Figure 1D).

It was previously reported (Lybecker *et al*., 2010) that recombinant BB0268 binds *in vitro* transcribed *B. burgdorferi* SR0440, a possible *B. burgdorferi* base-pairing sRNA denoted DsrA (Lybecker & Samuels, 2007) based on the well-characterized DsrA sRNA in *E. coli* (Majdalani *et al*., 1998). However, the electrophoretic mobility shift assays did not include negative controls and used high amounts of purified BB0268 (a maximum protein:RNA ratio of ∼823:1 molecules (Lybecker *et al*., 2010), compared to the maximum ∼15:1 ratio typically used for *E. coli* Hfq experiments (Zhang *et al*., 2002)). To test if BB0268 specifically interacts with SR0440, we carried out northern analysis on the *B. burgdorferi* BB0268-3XFLAG total, supernatant, and IP samples. In our experiment, no SR0440 was detected in the BB0268-3XFLAG IP sample (Figure S1D). In contrast, we observed a clear enrichment of *E. coli* DsrA in the *E. coli* Hfq-FLAG IP sample (Figure S1E). Collectively, these data indicate that BB0268 does not globally bind RNA or specifically bind a *B. burgdorferi* sRNA.

### BB0268 is not an Hfq homolog

A recent review conducted a systematic phyletic survey of Hfq homologs across 628 bacterial species representing all major phyla (Olejniczak & Storz, 2017). This identified an Hfq in two Leptospiraceae, *Leptospira* and *Turneriella*, but not in Borreliae. To ensure that no divergent version had been missed, we carried out a systematic search for Hfq homologs using sensitive sequence profile searches. These detected not only diverse bacterial Hfq proteins but also significantly identified the divergent archaeo-eukaryotic Sm clade proteins. Phylogenetic and divergence rate analysis of the prokaryotic Hfq proteins revealed that they fell into two broad clades: (i) the classic Hfq (Figure S2A); and (ii) the mobile Hfq (Figure S2B) clades. The classic Hfq clade is typified by the *E. coli* and *B. subtilis* Hfq proteins and is found in several major bacterial lineages, including Spirochaetes, such as Leptospiraceae. In the classic clade, *hfq* has gene-neighbors that are conserved across distant phyla including *miaA* (a tRNA modification gene), *hflX/hflK/hflC* (genes involved in ribosome release and as protein stability factors functioning with FtsH) and *mutL/mutS* (genes involved in DNA repair/ribosome rescue and release). The mobile Hfq clade is the only type of Hfq found in cyanobacteria and has been widely disseminated through lateral gene transfer across bacteria from an archaeal progenitor, probably through the medium of bacteriophages and plasmids, some of which also encode the protein. Sequence profiles for both these clades generated no statistically significant hits from the *B. burgdorferi* proteome establishing beyond a doubt that Hfq is absent in this organism. Further, the locus encoding *miaA* (*bb0821*; Figure S2C), showed no gene in the position usually occupied by Hfq homologs.

The lack of RNA-binding activity for BB0268 and our conservation analysis were consistent that Hfq is missing from *B. burgdorferi*, therefore, we looked further into the original *bb0268* study which reported BB0268 as an Hfq homolog (Lybecker *et al*., 2010). The study predicted that *B. burgdorferi* BB0268 has a protein secondary structure similar to *E. coli* and *S. aureus* Hfq. However, this is incompatible with AlphaFold2 secondary structure predictions (Jumper *et al*., 2021) for monomers of *E. coli* Hfq (Figure S3A) and BB0268 (Figure S3B) which showed no similarity between the two proteins. Hfq harbors an SH3-like β-sheet fold secondary structure (reviewed in (Brennan & Link, 2007)). In contrast, we predict that BB0268 harbors two transmembrane helices followed by an unstructured cytoplasmic tail (discussed below).

It was also previously reported that BB0268 complements an *E. coli* Δ*hfq* strain (Lybecker *et al*., 2010). Thus, we examined the effects of producing BB0268 in *E. coli*. We performed a growth curve using *E. coli* Δ*hfq* cells, but Δ*hfq* cells producing BB0268 grew worse than WT and BB0268 did not complement the *hfq*-dependent growth defect (Figure S4A). The levels of RpoS, the general stress-responsive σ factor (reviewed in (Gottesman, 2019)) are known to decrease in an *E. coli* Δ*hfq* strain (Muffler *et al*., 1996). It was previously proposed that BB0268 restored levels of RpoS, as measured by β-galactosidase activity from an *rpoS-lacZ* fusion in an *E. coli* Δ*hfq* background (Lybecker *et al*., 2010). To repeat this experiment, we examined *E. coli* RpoS levels using antibodies to the native protein (Figure S4B). RpoS decreased in Δ*hfq* samples at log and stationary phase, as expected, but we did not observe restoration of *E. coli* RpoS levels upon expression of *bb0268*. BB0268 production did lead to a modest RpoS increase in the stationary phase samples; however, it is unknown if this effect is direct or caused from stress responses invoked by BB0268 synthesis in *E. coli*.

The major role of *E. coli* Hfq is to facilitate RNA–RNA interactions and stabilize sRNAs, preventing them from degradation by RNases (reviewed in (Updegrove *et al*., 2016, Santiago-Frangos & Woodson, 2018)), but complementation of these *E. coli* Hfq phenotypes by *bb0268* was never tested. We isolated RNA from the same cultures used for immunoblot analysis (Figure S4B) and performed a northern analysis to measure the levels of various sRNAs (Figure S4C). We probed for the sRNA ChiX, one of the most enriched sRNAs bound by *E. coli* Hfq during growth in LB and highly unstable in an Δ*hfq* background (Melamed *et al*., 2020). No complementation was observed for ChiX levels in the Δ*hfq* cells expressing *bb0268*. We also examined the levels of two RpoS-dependent sRNAs, DsrA and SdsR (Majdalani *et al*., 1998, Fröhlich *et al*., 2012). Expression of *bb0268* in the Δ*hfq* strain failed to complement the effect of Δ*hfq* on sRNA levels in the log phase samples. At stationary phase, DsrA and SdsR bands were detected, albeit at negligible levels compared to the WT and the *hfq* complementation samples at the same time point. Collectively, these data demonstrated that ectopic expression of *bb0268* in *E. coli* does not significantly complement the effects of an *hfq* deletion mutant and the function of BB0268 is disparate from that of Hfq.

### *bb0268* is co-conserved with flagellar genes and σ^28^ across bacteria

To understand the true function of BB0268, we reexamined the genomic context and RNA expression of *bb0268*. The gene is embedded within the “*flgB* operon” and expression of the entire operon has been suggested to be constitutive, driven by a consensus σ^70^ promoter upstream of *flgB* (Ge *et al*., 1997b, Zhang *et al*., 2020). However, *B. burgdorferi* transcriptome mapping studies (Adams *et al*., 2017, Petroni *et al*., 2023) documented three predominant transcription start sites (TSSs), several 5′ processed ends, and several 3′ ends throughout the operon (Figure S5A). Therefore, this genomic region likely encodes multiple RNA products.

Northern analysis with a probe internal to *bb0268* (pink asterisk; Figure S5A) indicated that the gene is expressed in multiple transcripts across growth ranging in length from ∼500 to 6,000 nt (Figure S5B). No *bb0268* transcripts were detected by northern analysis using RNA isolated from a *B. burgdorferi* strain in which the entire *bb0268* ORF (480 nt) was replaced with a streptomycin (*aadA*) resistance cassette (1159 nt). Additional northern analysis with probes internal to the *bb0268*-neighboring genes *fapA* (*bb0267*), *flhG* (*bb0269*), and *flhB* (*bb0272*) (black asterisks; S5A and S5B) also detected multiple transcripts including a ∼6,000 nt band. Some transcripts detected with the *fapA* (*bb0267*) probe increased in the *bb0268* deletion samples, likely from the presence of the promoter for the *aadA* resistance cassette, which is oriented in the same direction of the operon. Together, these data document that the operon is expressed as multi- and single-gene products, with *bb0268* co-transcribed with *bb0272*– *bb0264*.

We next examined the conservation of BB0268 across bacteria by performing iterative and transitive sequence-profile and Hidden Markov Model searches (see Materials and Methods). These searches recovered a widespread family of BB0268 homologs with significant e-values (e < 10^-5^) from spirochetes, the PVC (Planctomyces, Verrucomicrobia, Chlamydia, Phycisphaerae, Omnitrophota) clade, flagellated members of the firmicute phylum (Bacillota), Nitrospinota, a subset of motile Chloroflexi, and certain other poorly studied bacterial clades (Desantisbacteria, Delongbacteria, Margulisbacteria, Glassbacteria, Ozemobacteria, Poribacteria). An analysis of the multiple sequence alignment and the predicted structure of these proteins (Figure S6A) revealed a two transmembrane (2TM) architecture with a well-conserved intramembrane arginine residue in the second transmembrane helix followed by a disordered, highly polar cytoplasmic tail of variable length culminating in a short, conserved C-terminal peptide with multiple hydrophobic amino acids and a basic residue.

Despite its broad phyletic distribution, the *bb0268* family has not been well characterized. Hence, we conducted a systematic analysis of the gene neighborhoods of this family to infer potential functional connections (Figure 2; predicted protein domains labeled for each gene). Without exceptions, all members of this superfamily are located within the flagellar superoperon across their broad phyletic distribution. Within this superoperon, the *bb0268* gene occurs at the junction between two genetic sub-modules for flagellar biosynthesis. First, there is a persistent association with the sub-module encoding two flagellar assembly proteins, FlhF (a flagellar GTPase), which regulates flagella number and structural organization and FlhG (FleN/MotR; a flagellar MinD NTPase homolog) which similarly has been implicated in the control of flagella count. Second, the *bb0268* gene associates with a sub-module encoding the flagellar σ factor, σ^28^ (Fl-Sigma; lost in *Borrelia*), FapA (flagellar assembly protein A), YjfB (an uncharacterized small protein), and a flagellar-associated transmembrane protein with a helix-turn-helix domain (TM+HTH). These observations strongly implicated *bb0268* in flagellar function.

**Figure 2.**
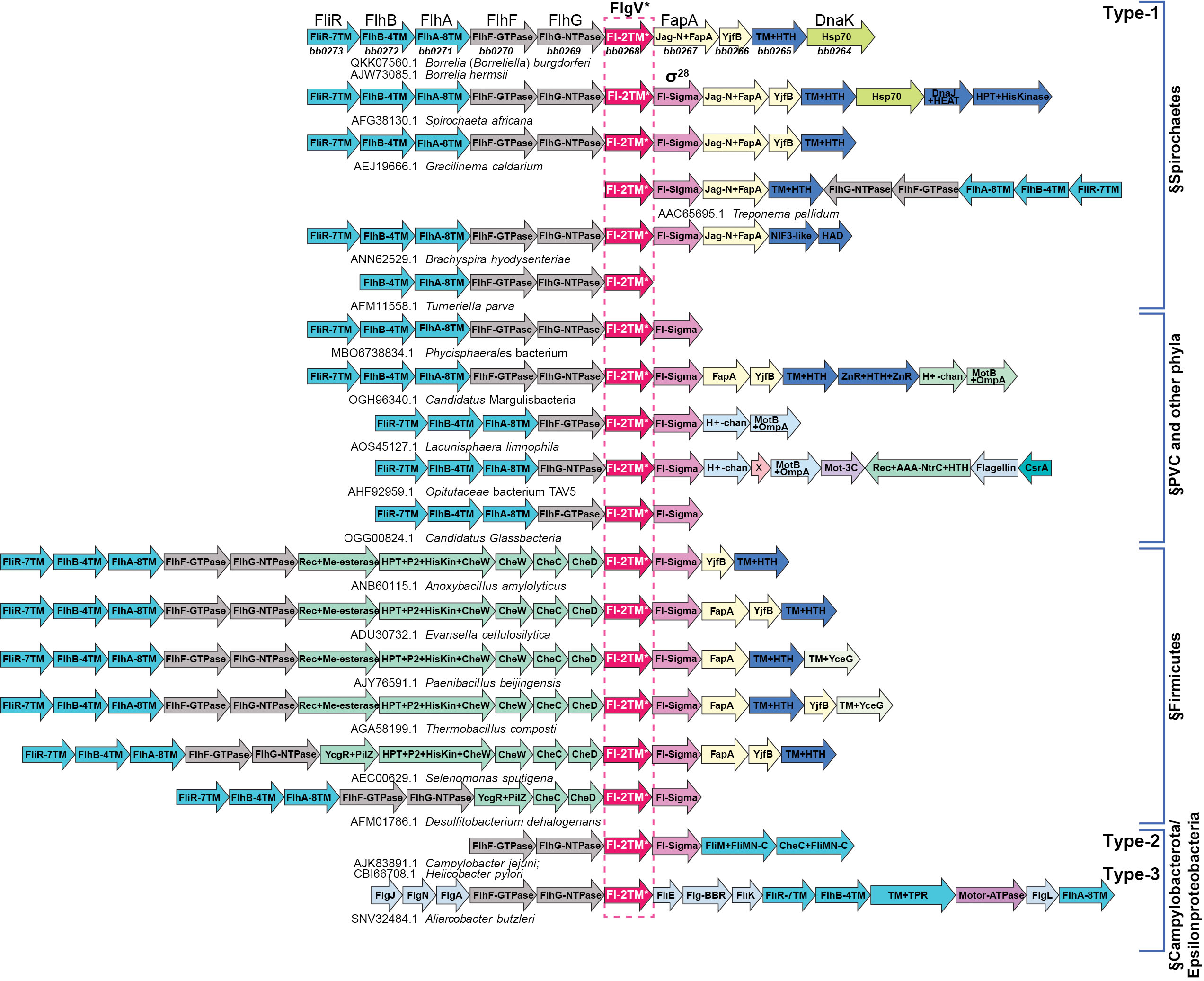
*bb0268* (*flgV*) is broadly conserved with flagellar genes and σ^28^ in bacteria. Examples of the flagellar superoperons from diverse bacterial lineages mentioned in the text are depicted, centered on the gene encoding FlgV, marked with a dashed box. The genes are shown as box arrows (only drawn to approximate size) and labeled with the domain architectures of the encoded proteins. The genes belonging to distinct genetic/functional submodules are distinguished using different colors. Each operon is labeled, below the box arrows, based on the GenBank accession of the anchor *flgV* gene and the species name. The encoded protein names from model systems are shown above the first row, and the corresponding *Borrelia* (*Borreliella*) gene names are listed below the operon from that organism. The versions are labeled as Type-1, -2, and -3 based on the type of FlgV encoded in the operon. All versions are Type-1 except the last two exemplars. The *Turneriella parva* superoperon is fragmented into more than one gene cluster, and the one containing *flgV* ends with that gene.

Interestingly, we also recovered homologs of a flagellar protein previously identified in the Campylobacterales (Campylobacter-Helicobacter) clade, named *flgV*, for its association with a motility phenotype in a *Campylobacter jejuni* transposon mutagenesis screen (Gao *et al*., 2014). Structural analysis of the Campylobacterales FlgV proteins revealed a 2TM architecture with a C-terminal cytoplasmic disordered tail congruent to the BB0268 family, though at lower significance (e-value = 0.1). Importantly, genome-context analysis revealed that Campylobacterales FlgV proteins too were encoded in the flagellar operons associated with the same two flagellar genetic sub-modules as the *bona fide bb0268* homologs (Figure 2). Taken together, these observations indicated the presence of a structurally similar 2TM protein in numerous bacteria that come in three distinct types (Figure 2): (i) Type-1, which is the most widely distributed, is defined by the above-mentioned BB0268-related proteins and are encoded in flagellar superoperons; (ii) Type-2, which includes the FlgV of Campylobacteria and Helicobacteria, typically are encoded in a short operon with just six flagellar genes; (iii) Type-3, which occur in Arcobacteria-like Campylobacterota and are encoded in the flagellar superoperon, but with a gene arrangement that is distinct from the first two types. Given its structural and gene-neighborhood equivalence, we renamed *B. burgdorferi bb0268* as *flgV,* keeping with the *Campylobacter jejuni* nomenclature. Despite being a widely distributed gene with a predicted flagellar function, *flgV* is absent in well-studied models such as *B. subtilis* and *E. coli*.

The *flgV* gene is frequently adjacent to the gene encoding σ^28^, the critical regulator of flagellar filamentation across bacteria. While all Spirochaetes encode *flgV*, it is striking that *Borrelia* are unique in that they specifically lack σ^28^ (Figure 2; *Turneriella* encodes σ^28^ at a different location from *flgV*). Thus, how *Borrelia* sp. regulate late-stage assembly of flagella has remained elusive. Here we sought to test the hypothesis that *flgV* impacts motility and flagellar assembly in *B. burgdorferi*.

### FlgV levels impact cell division and motility

To characterize *B. burgdorferi flgV*, we analyzed cells lacking and overexpressing the FlgV protein. To complement *flgV* in the deletion mutant, we altered the standard pBSV2G *B. burgdorferi* shuttle vector (Elias *et al*., 2003) to have the IPTG-inducible lactose promoter and the *lacI* gene codon optimized for *B. burgdorferi* (Blevins *et al*., 2007) (p_ind_). This resulted in a set of isogenic strains: WT (WT/p_ind_), *flgV* deletion (Δ*flgV*/p_ind_), and *flgV* complement (Δ*flgV*/p_ind_+*flgV*). To generate a *flgV* overexpression strain (++*flgV*) we cloned the *flgV* ORF sequence downstream of the constitutively active *flaB* promoter, into the standard pBSV2G vector in WT *B. burgdorferi* (WT/p_con_++*flgV*). WT spirochetes harboring the pBSV2G empty vector (WT/p) were used as a control for the ++*flgV* strain. We analyzed FlgV levels in these strains by immunoblot analysis, using antibodies to the native protein. FlgV was absent from the deletion strain and detected in the Δ*flgV*/p_ind_+*flgV* strain, albeit at slightly higher levels compared to WT (Figure 3A). Significantly higher FlgV levels were observed with the ++*flgV* construct (Figure 3B).

**Figure 3.**
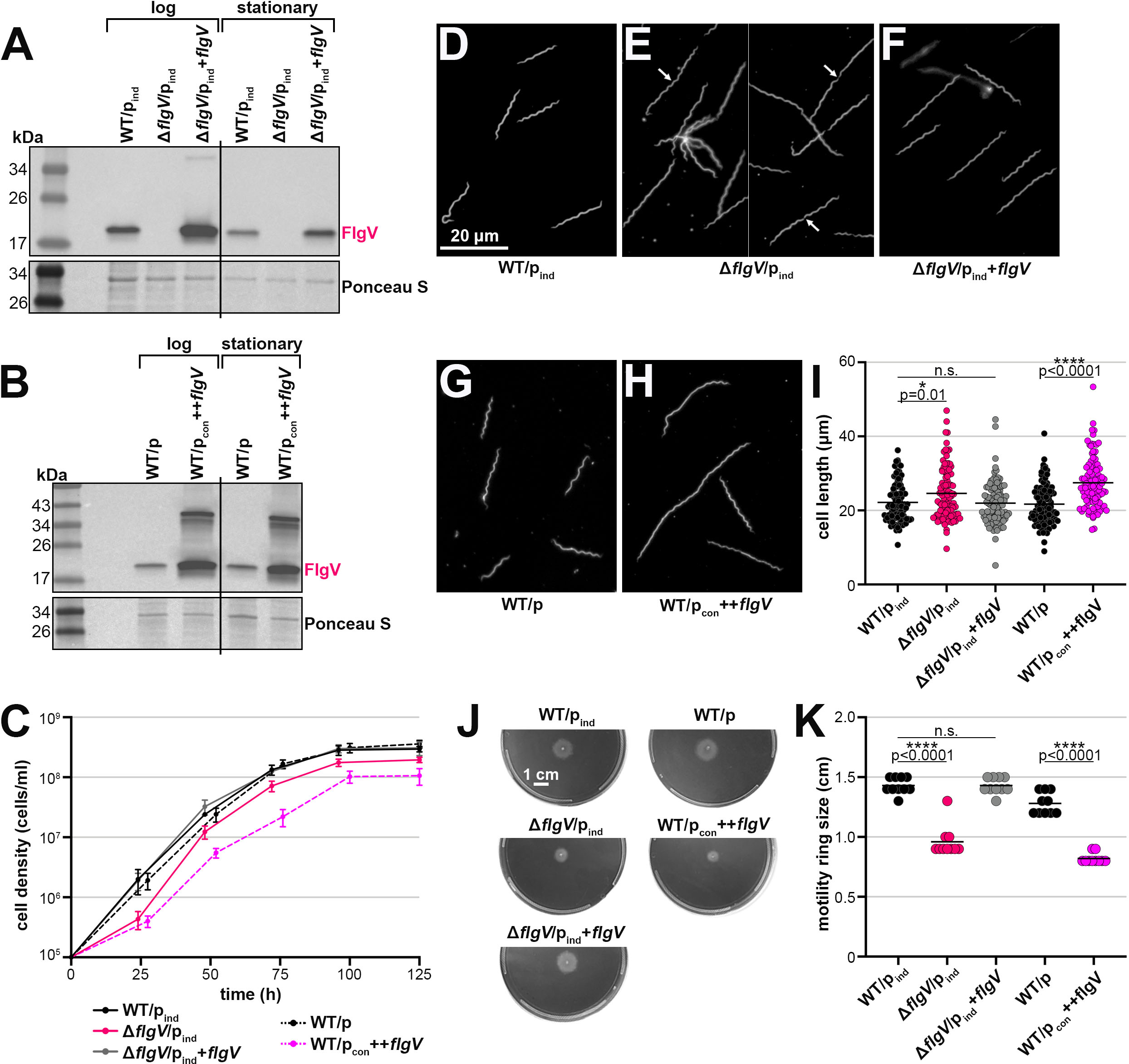
FlgV levels impact *B. burgdorferi* cell division and motility. (**A)** Immunoblot analysis of FlgV levels in WT, *flgV* deletion and *flgV* complementation strains. WT/p_ind_ (PA273), Δ*flgV*/p_ind_ (PA310), and Δ*flgV*/p_ind_+*flgV* (PA312) were grown with 0.1 mM IPTG, after dilution of the starter culture, to an average density of ∼1.1×10^7^ cells/ml (log) and ∼2.3×10^8^ cells/ml (stationary) and total protein was isolated. Δ*flgV*/p_ind_ (PA310) samples were collected 9 h after the WT/p_ind_ (PA273) and Δ*flgV*/p_ind_+*flgV* (PA312) samples, so that all cultures were collected at the same cell density. Protein extracts were separated on a Tris-Glycine gel, transferred to a membrane, stained with Ponceau S as a loading control, and probed with α-FlgV antibodies. Panel A was cropped to remove *B. burgdorferi* samples expressing *E. coli* Hfq. All samples are depicted in Figure S4E. Size markers are indicated. **(B)** Immunoblot analysis of FlgV levels in WT and *flgV* overexpression strains. WT/p (PA023) and WT/p_con_++*flgV* (PA267) were grown to an average density of ∼2.6×10^7^ cells/ml (log) and ∼1.8×10^8^ cells/ml (stationary) and total protein isolated. WT/p_con_++*flgV* (PA267) samples were collected 6 h after the WT/p (PA023) sample, so that all cultures were collected at the same cell density. Immunoblot was conducted as in panel A. **(C)** Growth curve of *B. burgdorferi* expressing different levels of FlgV. Cell growth was monitored by dark field microscopy enumeration at the indicated time points, after dilution of the starter culture to 1×10^5^ cells/ml. Each data point (circles) represents the mean of three biological replicates with the standard deviation. (**D–H**) Dark field microscopy images of representative (**D**) WT/p_ind_ (PA273), (**E**) Δ*flgV*/p_ind_ (PA310), (**F**) Δ*flgV*/p_ind_+*flgV* (PA312), (**G**) WT/p (PA023), and (**H**) WT/p_con_++*flgV* (PA267). All cultures were grown to an average density of ∼2.1×10^7^ cells/ml, washed with 1X PBS and imaged. For WT/p_ind_, Δ*flgV*/p_ind_, and Δ*flgV*/p_ind_+*flgV*, 0.1 mM IPTG was added at the subculture. White arrow indicates septa. Scale bar on panel D applies to panels D-H. **(I)** Quantification of spirochete length, for the strains in panels D–H. Approximately 100 cells were traced using the curve (spline) tool with ZEN 3.4 (blue edition) software. Each data point (circles) represents the length of one spirochete; the line corresponds to the mean length for each strain. Average length across WT/p_ind_, Δ*flgV*/p_ind_, and Δ*flgV*/p_ind_+*flgV* samples or WT/p and WT/p_con_++*flgV* were compared by one-way ANOVA with Tukey’s multiple comparisons test or t test, respectively, GraphPad Prism 9.5.1 (n.s., not significant). **(J)** Motility assay of *B. burgdorferi* expressing different levels of FlgV. Representative plates of each strain were photographed 9 d after inoculating into a BSKII 0.35% agarose plate. Scale bar on first plate applies to all plates. **(K)** Quantification of spirochete motility, for the strains in panel J. Each data point (circles) represents the motility ring size (distance spirochetes spread from inoculation site) for one sample; the line corresponds to the mean motility ring size for each strain. The statistical analysis was performed as reported for panel I.

We first measured the growth of the *flgV* strains by dark-field microscopy enumeration (Figure 3C). When *flgV* was deleted, the spirochetes had a moderate *flgV*-dependent lag in cell growth. Complementation by IPTG-induction of *flgV* resulted in full restoration of growth compared to the Δ*flgV*/p_ind_ strain. FlgV overproduction also negatively impacted cell growth. There was no difference in growth between the two WT control strains.

We observed differences in cell morphology when *flgV* levels were altered. Spirochetes grown to exponential phase and lacking *flgV* were longer than WT *B. burgdorferi* (Figure 3D and 3E). Closer inspection of the longer Δ*flgV*/p_ind_ cells showed conjoined Δ*flgV* spirochetes and the presence of septa (white arrows; Figure 3E). The *flgV*-dependent morphology phenotypes were complemented with p_ind_+*flgV* (Figure 3F). Overproduction of FlgV also resulted in longer spirochetes, compared to WT cells (Figure 3G and 3H). To quantify these observations, we measured the length of ∼100 cells per strain using dark-field microscopy (Figure 3I). Both the lack of and elevated levels of FlgV significantly increased the average spirochete length in exponential phase. There was no difference in the cell length of Δ*flgV*/p_ind_ or WT/p_con_++*flgV B. burgdorferi* in stationary phase, compared to the WT controls (Figure S4F).

Cell division and motility are likely inherently linked in spirochetes, as efficient separation at the mid-cell septum requires functional flagella (Lin *et al*., 2015). To measure the contribution of *flgV* to spirochete motility, we inoculated the *flgV* strains into semisolid plates (Figure 3J) and measured the diameter of motility rings (Figure 3K). On average, Δ*flgV/*p_ind_ spirochetes formed smaller rings compared to WT spirochetes. The deletion phenotype was rescued by *flgV* complementation. A similar defect in motility was observed with spirochetes overproducing FlgV. Collectively our data indicate levels of FlgV must be tightly regulated, as both too much or too little protein resulted in cell division and motility defects.

### FlgV is membrane associated and localized to the cell poles and mid-cell

Members of the FlgV protein family (Figure S6A) have a similar structure of two predicted N-terminal transmembrane helices (amino acids 13 to 35 and 40 to 57, in *B. burgdorferi*). To experimentally determine FlgV localization, total cell lysates from WT *B. burgdorferi* were fractionated into soluble and membrane portions and analyzed by immunoblot analysis (Figure 4A). FlgV was restricted to the membrane fraction like membrane-associated outer surface protein C (OspC) and not like soluble superoxide dismutase A (SodA). Previous gel filtration chromatography reported that FlgV likely forms a dimer *in vitro* (Lybecker *et al*., 2010), consistent with this, we observed a higher molecular weight FlgV band (Figure 4A) that was more apparent in our co-IP (Figure 1A) and overexpression (Figure 3B) studies.

**Figure 4.**
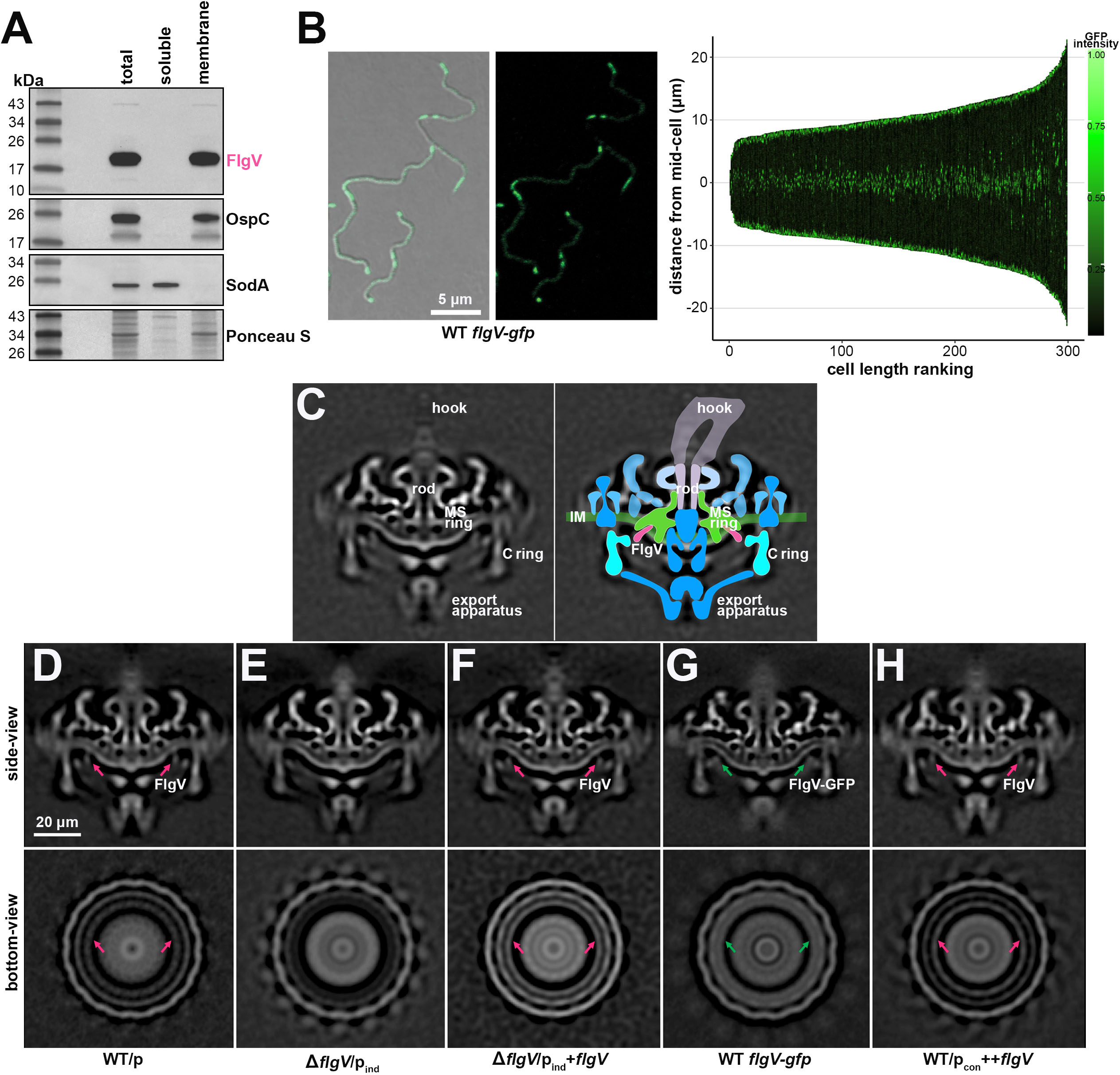
FlgV is localized to the flagellar motor. **(A)** Immunoblot analysis of *B. burgdorferi* total protein lysate fractionated into soluble and membrane samples. WT *B. burgdorferi* (PA001) were grown to a density of ∼2.2×10^7^ cells/ml and then lysed by sonication, total protein was isolated, and the samples were fractioned by ultracentrifugation. Protein extracts were separated on a Tris-Glycine gel, transferred to a membrane, stained with Ponceau S as a loading control, and probed with α-FlgV, α-OspC, and α-SodA antibodies. Proteins were probed sequentially on the same membrane; size markers are indicated. **(B)** Localization of FlgV-GFP by confocal microscopy. *B. burgdorferi* (PA402) were grown to a density of ∼2.0×10^7^ cells/ml, prior to imaging. Left panels: Bright field and fluorescence composite (left micrograph) and fluorescence-only (right micrograph) are shown. Scale bar on left micrograph also applies to right micrograph. Right panel: Demograph of the GFP profile for 300 spirochetes, cells were positioned in order of increasing length. **(C)** Reconstructed cryo-ET central (side-view) cross section of WT/p (PA023) *B. burgdorferi* flagellar motor (left) with overlaid cartoon model of the flagellar motor (right). The inner membrane (IM, green), export apparatus (blue), C ring (light blue), MS ring (green), rod (lavender), hook (light purple) and FlgV (dark pink) are labeled. (**D–H**) Reconstructed central (side-view) and bottom-view cryo-ET cross sections of (**D**) WT/p (PA023; repeated from panel C), (**E**) Δ*flgV*/p_ind_ (PA310), (**F**) Δ*flgV*/p_ind_+*flgV* (PA312), (**G**) WT *flgV-gfp* (PA402), and (**H**) WT/p_con_++*flgV* (PA267) *B. burgdorferi*. All cultures were grown to an average density of ∼3.6×10^7^ cells/ml, washed with 1X PBS, mixed with 10 nm gold particles, deposited on glow-discharged grids and imaged. Tomographs were generated, aligned and reconstructed for each strain. Densities predicted to correlate with FlgV (pink arrow) and FlgV-GFP (green arrow) are indicated. Scale bar on panel D applies to panels D–H.

We created a FlgV C-terminal fusion to codon-optimized green fluorescent protein (Takacs *et al*., 2018) at the native *flgV* locus (Figure S6B). Spirochetes producing FlgV-GFP were grown to logarithmic phase and imaged by fluorescence confocal microscopy (Figure 4B). We constructed a demograph where the GFP profile for 300 spirochetes was displayed for each cell, by aligning spirochetes in order of increasing length. FlgV-GFP localized to the cell poles, which overlaps with the location of the flagellar motors (Liu *et al*., 2009). The analysis also revealed GFP intensity at two puncta in the middle of each spirochete. The mid-cell coincides with the point of septation of a dividing spirochete, thus the new poles of a future daughter cell (Jutras *et al*., 2016) and location of future flagella. In the longest spirochetes we examined, which may actually be two spirochetes about to divide, we sometimes observed two internal doublet-puncta of GFP intensity (Figure 4B; demograph), which could correspond to the future mid-cell of the two new spirochetes. Similar FlgV-GFP localization was observed for cells grown to stationary phase (Figure S6C). Together, our observations support a hypothesis that membrane-associated FlgV co-localizes with functional flagellar motors at the cell ends and nascent flagella motors at the mid-cell.

### FlgV is a component of the flagellar motor

To investigate a possible FlgV association with the flagellar motor, we analyzed *in situ* structures of flagellar motors using cryo-electron tomography (cryo-ET). The *in situ* motor structure from WT/p cells revealed major components of the flagellar basal body including the export apparatus, rotor complex (C and MS rings), stator complex, and rod (Figure 4C and 4D), as previously defined (Liu *et al*., 2009). We also determined *in situ* structures of the flagellar motors from Δ*flgV*/p_ind_ (Figure 4E) and Δ*flgV*/p_ind_+*flgV* (Figure 4F) *B. burgdorferi*. The structures of the Δ*flgV* motor exhibited many of the same features as the WT motors; however, Δ*flgV* motors lacked a density adjacent to the C and MS rings of the rotor (pink arrows; Figure 4D and 4E). Complementation restored the absent cryo-ET densities in the Δ*flgV* spirochete motors (Figure 4F). To further demonstrate the association of this density with FlgV, we analyzed the motors of *flgV-gfp* spirochetes. The FlgV C-terminal fusion to GFP increases the size of FlgV by 237 aa. We observed a significant increase in the FlgV-associated cryo-ET density for WT *flgV-gfp* samples (green arrows; Figure 4G). These data support the model that FlgV localizes to the flagellar motor between the C and MS rings of the flagellar rotor (Figure 4C), and the corresponding cryo-ET density is not part of the C-ring protein FliG2 as suggested previously (Liu *et al*., 2009). We also determined the motor structure when FlgV was overproduced (Figure 4H). Surprisingly, we observed no significant difference in the WT/p_con_++*flgV* motor structures compared to those in WT/p, suggesting that FlgV overproduction did not alter the flagellar motor structure.

### FlgV impacts the assembly of flagellar filaments

*B. burgdorferi* B31 standardly has 7–11 flagellar motors at each cell pole, and each flagellar motor is associated with one hook and a long filament that extends through the periplasmic space (Hovind-Hougen, 1984, Qin *et al*., 2018). To understand the cell division and motility defects associated with altering FlgV levels, we used cryo-ET to visualize periplasmic flagella when *flgV* was deleted or the protein was overexpressed. As expected, in WT/p spirochetes we observed 7–11 flagellar filaments that formed a ribbon-like structure along the cell body (Figure 5A; representative cell with 9 flagellar filaments). Δ*flgV*/p_ind_ spirochetes had similar, but fewer, ribbon-like flagellar filaments, suggesting that the flagellar filaments are slightly shorter or reduced in number (Figure 5B; representative cell with 7 flagellar filaments). *flgV* complementation restored the number of filaments to WT/p levels (Figure 5C; representative cell with 9 flagellar filaments). Strikingly, when FlgV was overproduced fewer ribbon-like structures were observed, suggesting that the flagellar filaments are significantly shorter or reduced in number (Figure 5D; representative cell with 4 flagellar filaments).

**Figure 5.**
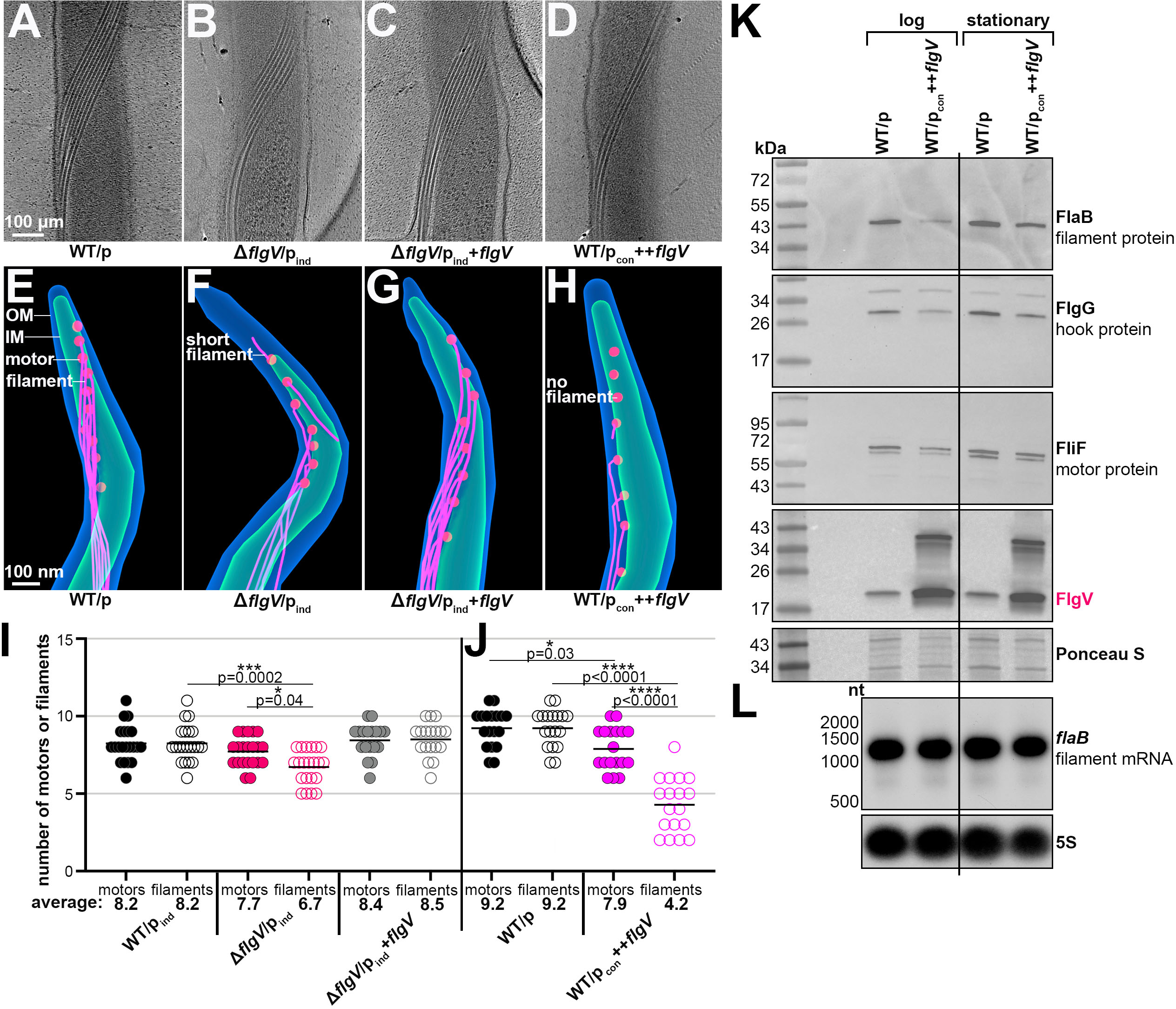
FlgV levels impact the assembly of flagellar filaments. (**A–D**) surface-view cryo-ET micrograph of representative central periplasmic cell sections of (**A**) WT/p (PA023), (**B**) Δ*flgV*/p_ind_ (PA310), (**C**) Δ*flgV*/p_ind_+*flgV* (PA312), and (**D**) WT/p_con_++*flgV* (PA267) *B. burgdorferi*. All cultures were grown to an average density of ∼3.6×10^7^ cells/ml, washed with 1X PBS, mixed with 10 nm gold particles, deposited on glow-discharged grids, and imaged. Scale bar on panel A applies to panels A–D. (**E**–**H**) Projection images of a representative cell pole for each strain, described in panels A–D. The outer membrane (blue), inner membrane (green), flagellar motors (red) and flagellar hooks and filaments (pink), were manually segmented in IMOD and are indicated. Scale bar on panel E applies to panels E–H. See Supplementary Videos S1-S4. (**I–J**) Quantification of flagellar motors and filaments. Individual spirochetes were tallied for number of motors and filaments at one cell pole. Each data point (circles) represents one spirochete; the line corresponds to the mean number of motors or filaments for each strain. Panels I and J were independent experiments; panel I: 0.1 mM IPTG was added at the subculture and WT/p_ind_ (PA273), Δ*flgV*/p_ind_ (PA310), and Δ*flgV*/p_ind_+*flgV* (PA312) cultures were grown to an average density of 3.8×10^6^ cells/ml; panel J: WT/p (PA023) and WT/p_con_++*flgV* (PA267) were grown to an average density of 2.3×10^7^ cells/ml. Average motor and filament numbers per sample were compared by one-way ANOVA with Tukey’s multiple comparisons test, GraphPad Prism 9.5.1. (**K**) Immunoblot analysis of flagellar structural proteins when FlgV is overexpressed. The membrane from Figure 3B was stripped and reprobed with α-FlaB (Barbour *et al*., 1986), α-FlgG and α-FliF antibodies. The FlgV panel was repeated from Figure 3B; the Ponceau S panel is the same as Figure 3B, but shows a different region; size markers are indicated. (**L**) Northern analysis of *flaB* levels. Total RNA was isolated from a portion of the same cultures used in panel K. RNA was separated on an agarose gel, transferred to a membrane and probed for *flaB*. The membrane was stripped and probed for 5S as a loading control; size markers are indicated.

To further explore and quantify our observations, we used cryo-ET to examine the spirochete cell poles and enumerate both flagellar motors and the associated filaments (Figure 5E–H; representative tomograph projections for each strain; Videos S1-S4). In WT/p spirochetes, we observed equal numbers of motors and filaments at the cell poles (Figure 5E). In contrast, Δ*flgV*/p_ind_ spirochetes had flagellar motors with incomplete, short flagellar filaments (Figure 5F) or motors that lacked a filament entirely. Complementation restored the filaments to WT/p levels (Figure 5G). In spirochetes overproducing FlgV, roughly half of flagellar motors lacked filaments or had incomplete, short flagellar filaments (Figure 5H). We observed no difference in the position of flagella across any *B. burgdorferi* strain. We counted the number of motors and filaments at one cell pole for ∼20 cells per strain. WT/p, Δ*flgV*/p_ind_, and Δ*flgV*/p_ind_+*flgV* cells all had similar numbers of flagellar motors (Figure 5I; mean values of 8.2 (n=18), 7.7 (n=21) and 8.4 (n=18), respectively). However, there was a significant reduction in the average number of flagellar filaments for Δ*flgV*/p_ind_, mean value of 6.7 (n=21), compared to WT/p, 8.2 (n=18) and the *flgV* complemented spirochetes, 8.5 (n=18), in which, the average number of motors and filaments were the same. In a separate experiment, we tallied the flagellar motors and filaments in spirochetes when *flgV* was overexpressed (Figure 5J). There were some variations in the numbers for WT/p_ind_ and WT/p *B. burgdorferi* between experiments (Figures 5I and 5J), possibly due to differences in the precise cell density of the sampled culture. Nevertheless, there consistently was a reduction in the average number of flagellar motors in *B. burgdorferi* overexpressing *flgV*, with a mean value of 7.9 (n=18), compared to WT/p spirochete motors, mean value of 9.2 (n=21). More strikingly, there was a significant reduction in the average number of flagellar filaments in spirochetes overproducing FlgV, mean value of 4.2 (n=18), lower than the number of flagellar motors in the overexpression strain and lower than the number of flagellar filaments in WT/p spirochetes, mean value of 9.2 (n=21). Collectively, our data provide direct evidence that changes in FlgV levels have profound impacts on the assembly of the flagellar filament, while they have limited impacts on the structure, number, and position of flagellar motors.

To further investigate the impact of *flgV* overexpression, we examined the total protein levels for specific components of the flagellar motor and filament. In *B. burgdorferi*, the flagellar filament is comprised of the major flagellin protein, FlaB, and a minor sheath protein, FlaA (Ge *et al*., 1998, Motaleb *et al*., 2004). Compared to WT/p samples, FlaB levels decreased in lysates isolated from WT/p_con_++*flgV B. burgdorferi* at both logarithmic and stationary phase (Figure 5K). We also observed a ++*flgV*-dependent decrease in FlgG, a hook protein which connects the flagellar filament to the motor. The levels of the motor protein FliF also decreased with *flgV* overexpression. These FlgV-effects were more prominent in the logarithmic phase samples, compared to the stationary phase samples. We also isolated and analyzed RNA from WT/p and WT/p_con_++*flgV B. burgdorferi* from the same experiment. Northern analysis indicated no change in the levels of *flaB* mRNA when *flgV* is overexpressed (Figure 5L). These data support a model that FlgV impacts flagellar assembly post-transcriptionally. We also examined FlaB levels when *flgV* was deleted; no significant decrease in FlaB protein (Figure S7A) or mRNA levels (Figure S7B) were observed. We performed a second, independent experiment with the *flgV* strains and arrived at the same conclusions (Figure S7C).

Collectively, our findings implicate FlgV as an important component of the flagellar motor that impacts the late-stage assembly of flagellar filaments. Given this effect on the flagella, we hypothesized that FlgV could play a role in controlling spirochete motility during the tick– mammal enzootic cycle.

### *flgV* is dispensable for *B. burgdorferi* survival and replication in ticks

*B. burgdorferi* have been observed to be non-motile in the midguts of infected unfed ticks but become motile during the bloodmeal (Dunham-Ems *et al*., 2009). To determine the importance of *flgV* throughout the enzootic cycle of *B. burgdorferi*, we artificially infected larval ticks by submerging pools of naïve *Ixodes scapularis* ticks in BSKII medium (Policastro & Schwan, 2003) containing WT/p, Δ*flgV*/p or WT/p_con_++*flgV* spirochetes and monitored the infection at each stage of tick-feeding on mice (Figure 6A). A subset of artificially-infected, unfed larval ticks were assayed for *B. burgdorferi* burden by crushing and plating for colony-forming units (CFUs) in solid BSKII medium (Grimm *et al*., 2005). All strains colonized the unfed larvae at the same level, following the immersion procedure, confirming successful artificial infection (Figure 6B).

**Figure 6.**
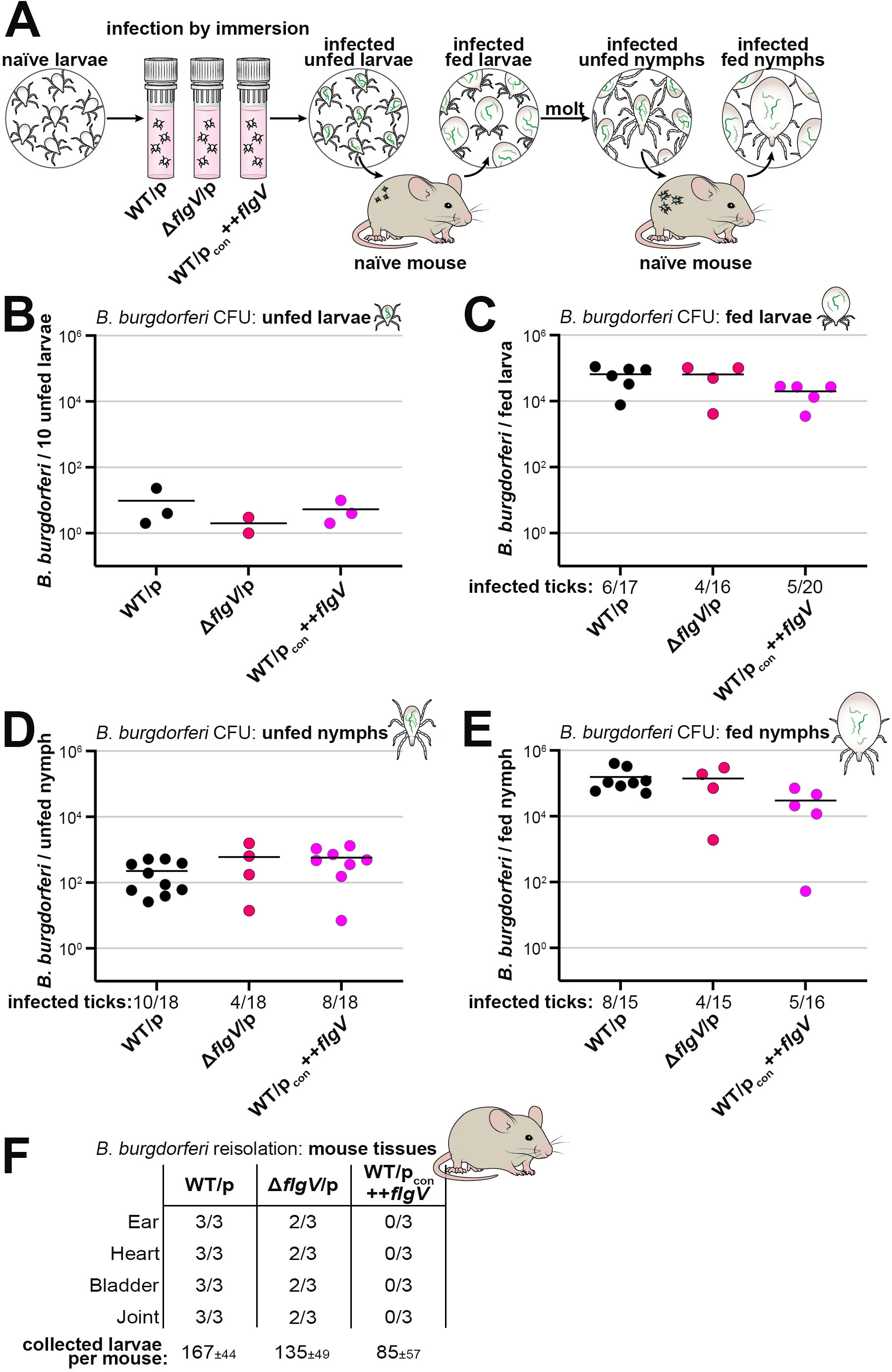
*flgV* is not important for *B. burgdorferi* replication or survival in *Ixodes* ticks. (**A**) Schematic of artificial tick infection by immersion used for tick-mouse infection study. Naïve, unfed larvae were desiccated and submerged in BSKII containing WT/p (PA023), Δ*flgV*/p (PA257), or WT/p_con_++*flgV* (PA267) *B. burgdorferi*. (**B–E**) Ticks were assayed for infectivity at each life-stage: (**B**) unfed larvae, (**C**) fed larvae, (**D**) unfed nymphs, and (**E**) fed nymphs. Ticks were surface sterilized, crushed and plated in solid BSKII containing RPA cocktail and gentamycin. Data points (circles) represent the number of colony-forming units (CFUs) for infected ticks. For unfed larvae (panel B), 3 groups of 10 unfed larvae from each strain were assayed for infectivity; one plate of the Δ*flgV*/p sample was contaminated and not able to be counted. Ticks from all other life-stages (panels C–E), were assayed individually for infectivity; the number of individual infected ticks over the number of individual crushed ticks is listed below the *x*-axis. The average numbers of *B. burgdorferi* per tick, for each tick life-stage, were compared by one-way ANOVA with Tukey’s multiple comparisons test, GraphPad Prism 9.5.1; no significant difference was observed for any tick life-stage. (**F**) Reisolation of *B. burgdorferi* from mouse tissues after larval tick feeding. 3 mice per group were assessed for infection 3 weeks post larval tick-feeding. Ear, heart, bladder and joint tissues were collected and analyzed for spirochete reisolation in BSKII by dark-field microscopy enumeration. Average numbers of larval ticks collected per mouse are indicated.

To test the ability of Δ*flgV* or ++*flgV* spirochetes to replicate in feeding ticks, infected larvae were fed to repletion (allowed to naturally fall off) on naïve mice and collected for analysis. The spirochete burden for individual fed larval ticks was determined by enumeration of CFUs. Not all ticks acquire *B. burgdorferi* by immersion infection (Policastro & Schwan, 2003), and therefore, it is important to quantify the initial infection efficiency for each bacterial strain. The larval tick-infection efficiency of WT *B. burgdorferi* was calculated at 35% (6 infection positive ticks/17 total assayed ticks), which was similar to larvae infected with Δ*flgV*/p (25%; 4/16 ticks) or WT/p_con_++*flgV* (25%; 5/20 ticks) (Figure 6C; reported below *x*-axis). We then quantified and compared the spirochete burdens for only the infected ticks in each group. There was no significant difference in the average number of *B. burgdorferi* per tick when *flgV* was deleted or FlgV was overproduced, compared to ticks infected with WT bacteria (Figure 6C).

Fed larvae were allowed to molt into nymphs and assessed for *B. burgdorferi* infection post-molt. There was no difference in spirochete survival through the molt for the *flgV* strains, compared to WT spirochetes (Figure 6D). Nymphs were fed to repletion on naïve mice and again, there was no significant difference in the average number of spirochetes per tick for any *B. burgdorferi* strain (Figure 6E). These data document that *flgV* is dispensable for *B. burgdorferi* survival and replication in larval and nymphal ticks.

We also determined the ability of the *B. burgdorferi* strains to infect mice by reisolating spirochetes from tissues approximately three weeks after the larval tick bite (Figure 6F). WT spirochetes were reisolated from the ear, heart, bladder and joint tissues of all three mice. In contrast, Δ*flgV*/p spirochetes were detected in all tissues of only two out of three mice and there was no reisolation of *flgV* overexpression spirochetes from any mouse tissue. In the nymphal tick feeding, we did not collect enough nymphs per mouse to confidently assess defects in mouse infectivity. Our data show that *flgV* is important for efficient *B. burgdorferi*-mammalian infection by tick bite and led us to further examine the consequences of altering *flgV* levels during mouse infection.

### *flgV* is critical for *B. burgdorferi* dissemination and infectivity in mice

Aspects of tick bite transmission of *B. burgdorferi* can be modeled by intradermal needle inoculation of mice (Figure 7A). Successful mammalian infection by *B. burgdorferi* begins with intradermal skin inoculation of spirochetes, which is followed by localized replication of bacteria at the inoculation site (Hodzic *et al*., 2002, Hodzic *et al*., 2003). At 5 to 7 days after the intradermal needle injection, spirochetes disseminate and are detectable in the bloodstream, with a peak in bacteriemia at day 6 (Aranjuez *et al*., 2019). At 7 to 10 days after inoculation, spirochetes exit the blood and start to be detectable in distal tissues, such as a secondary skin sites and ear, heart, and joint tissues, where long-term colonization is established (Hodzic *et al*., 2002, Hodzic *et al*., 2003).

**Figure 7.**
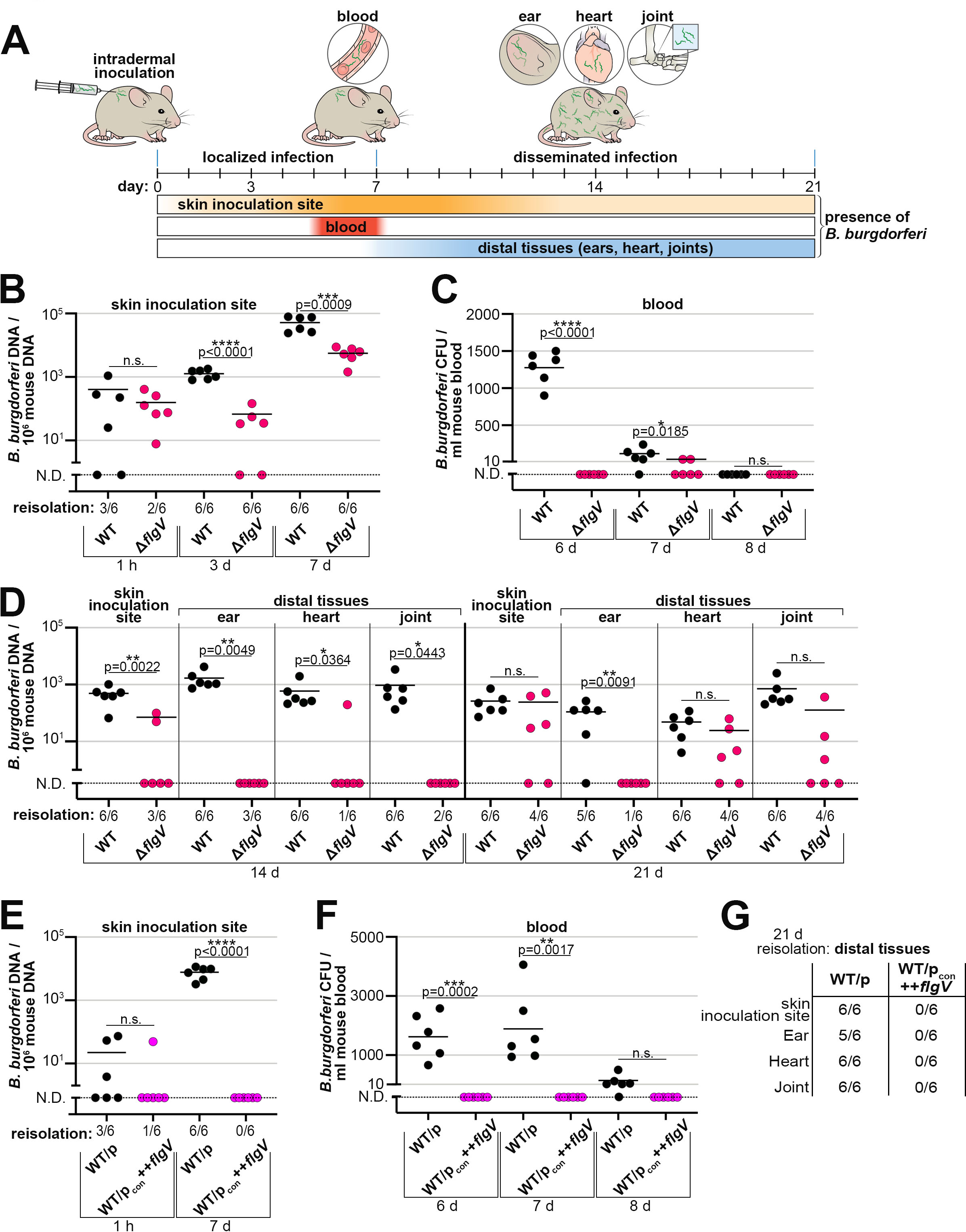
*flgV* is important for timely and productive dissemination in mice. (**A**) Schematic of mouse-*B. burgdorferi* infection kinetics by intradermal needle inoculation. (**B–D**) Mice were infected intradermally with 1×10^4^ WT (PA001) or Δ*flgV* (PA251) *B. burgdorferi*. After the indicated times, the (**B**) skin inoculation site, (**C**) blood, and (**D**) distal tissues were assessed for spirochete infectivity and burden. (**E–G**) In a separate experiment, mice were infected intradermally with 1×10^4^ WT/p (PA023) or WT/p_con_++*flgV* (PA267) *B. burgdorferi*. After the indicated times, the (**E**) skin inoculation site, (**F**) blood, and (**G**) distal tissues assessed for the presence of spirochetes. For panels B–G: Each data point (circles) represents one mouse; samples with no detectable spirochetes are indicated by a data point on the dotted line (N.D. = not detected). The mean for each strain is indicated by a line, only detected data were used to calculate the mean. For quantitative-PCR analysis, DNA was extracted from tissues and *B. burgdorferi* load was measured by quantifying *B. burgdorferi flaB* copies normalized to 10^6^ mouse *nid* copies, in technical triplicate for each sample. For bacteremia analysis, blood was collected, diluted in BSKII and plated. For tissue reisolation, skin inoculation site, ear, heart, and joint tissues were collected, submerged in BSKII and enumerated by dark-field microscopy. The calculated means between WT versus Δ*flgV* or WT/p versus WT/p_con_++*flgV* for each tissue/blood at a given timepoint were compared by a one-tailed, unpaired t test; for all statistical analyses N.D. data points were included as zero; GraphPad Prism 9.5.1; (n.s., not significant).

Using this model, we evaluated the contribution of *flgV* at each stage of mouse infection by quantifying spirochete loads and reisolating spirochetes from tissues. Groups of mice were intradermally inoculated with WT and Δ*flgV* spirochetes and infection parameters were assessed over a three-week infection time course. We first confirmed that equal number of WT and Δ*flgV* spirochetes were inoculated into the skin 1 hour after intradermal delivery by extracting DNA and quantifying the number of *B. burgdorferi* by qPCR (Figure 7B). There was no significant difference between WT or Δ*flgV* spirochete burden in the skin inoculation sites at the 1-hour timepoint. In parallel, using a section of the same skin sample examined by qPCR, we assayed the ability to reisolate WT or Δ*flgV* spirochetes from the inoculation site in liquid medium (Figure 7B; reported below the *x*-axis). Again, we observed no significant difference between the two strains. As expected, the numbers of WT spirochetes expanded in the skin inoculation site from 1 hour to 7 days post-infection. We were able to reisolate both WT and Δ*flgV* spirochetes from the inoculation site of all mice at the 3- and 7-day timepoints. However, compared to WT spirochetes, there were significantly fewer Δ*flgV* spirochetes in the skin inoculation sites at 3 and 7 days. Our data indicate that *flgV* is important for maximum *B. burgdorferi* expansion in the skin, during the first 7 days of infection.

Moving away from the site of injection, we examined the spirochete load in blood during hematogenous dissemination in mice by measuring spirochete bacteremia. Following an intradermal needle injection, WT *B. burgdorferi* are only detectable in blood approximately 5–7 days post-inoculation (Aranjuez *et al*., 2019). Blood was collected from mice at days 6, 7, and 8 post-injection and plated for *B. burgdorferi* CFUs (Figure 7C). As expected, WT *B. burgdorferi* were detected in blood at day 6, with a decrease in numbers at day 7 and below the limit of detection by day 8. Strikingly, no Δ*flgV* spirochetes were detected at day 6 and only two mice had detectable Δ*flgV* spirochetes in blood at day 7. Therefore, Δ*flgV* spirochetes may be delayed in entering the bloodstream, be at low numbers during hematogenous spread and/or be rapidly eliminated from the blood, ultimately attenuating spirochete dissemination to distal tissues.

We finally assessed the presence and amount of disseminated spirochetes in distal tissues. The initial skin inoculation site and distal tissues: ear, heart, and joint, were harvested and subjected to reisolation and qPCR analysis at 14 and 21 days post-inoculation (Figure 7D). At the 14-day timepoint, all tissues (24/24 of total skin, ear, heart and joint tissues) from the mice infected with WT *B. burgdorferi* were reisolation positive with quantifiable spirochetes. Only 37.5% (9/24) of mouse tissues were reisolation positive for Δ*flgV B. burgdorferi* at day 14 (Figure 7D; reported for each tissue below the *x*-axis). The numbers of Δ*flgV* spirochetes in tissues were generally below the limit of detection by qPCR at the 14-day timepoint. No Δ*flgV* spirochetes were detected in ear and joint tissues, and only one heart and two skin samples had detectable Δ*flgV* spirochetes. At 21 days-post inoculation, again WT spirochetes were reisolated and quantified from all tissues, excluding one ear sample (23/24). For the Δ*flgV B. burgdorferi* infection, 54.2% (13/24) of mouse skin, heart and joint samples were reisolation positive at day 21 (Figure 7D; reported below the *x*-axis). Despite low dissemination and infection rates, quantification of these tissues revealed no significant difference between the numbers of detected WT and Δ*flgV* spirochetes. These data suggest that if a tissue becomes infected with Δ*flgV B. burgdorferi*, the spirochetes ultimately can reach a similar level of tissue colonization, compared to WT *B. burgdorferi*. In contrast, and strikingly, only one mouse ear was reisolation positive for Δ*flgV* spirochetes and no Δ*flgV* spirochetes were detected, above the limit of detection, in any ear sample at day 21. Overall, we detected few Δ*flgV* spirochetes at 14 days post inoculation, but at 21 days post inoculation Δ*flgV* spirochetes were detected in a majority of mice. This suggests a *flgV*-dependent delay in the kinetics of *B. burgdorferi* dissemination and infection.

We also assayed mice that were intradermally inoculated with *B. burgdorferi* overproducing FlgV. Remarkably, no WT/p_con_++*flgV* spirochetes were detected in the skin inoculation sites by qPCR or tissue reisolation at 7 days post inoculation (Figure 7E). Spirochetes overproducing FlgV were also not detected in the blood at 6-, 7- or 8-days post needle injection (Figure 7F). Finally, no WT/p_con_++*flgV* spirochetes were able to be reisolated from any mouse tissue 21 days after the intradermal inoculation (Figure 7G). In contrast, WT/p spirochetes were robustly detected in the inoculation site (Figure 7E), in the blood (Figure 7F) and in distal tissues (Figure 7G). These data, consistent with our previous observations, indicate that higher levels of FlgV have greater consequences than *flgV* deletion on flagellar filamentation and *B. burgdorferi* infection of the mammalian host.

Collectively, our analysis of key timepoints of *B. burgdorferi* infection of mice documented that FlgV, and thus proper flagellar filamentation, is critical for timely and productive dissemination during an infection model of Lyme disease.

## DISCUSSION

Here we report a novel component of *B. burgdorferi* flagellar motors that impacts flagellar assembly, and thus significantly impacts motility and dissemination during mammalian infection. A previous study reported that *bb0268* (*flgV*) encodes an atypical Hfq protein (Lybecker *et al*., 2010). Thus, we set out to characterize the RNAs associated with BB0268, but in the absence of identifying any BB0268-bound RNAs, discovered *bb0268* was misannotated.

### BB0268 (FlgV) is not an Hfq homolog or an RNA-binding protein

RNA-binding proteins have been shown to play critical roles in modulating gene expression and are required for infection across a broad range of bacteria (reviewed in (Holmqvist & Vogel, 2018)). These RNA-binding proteins can facilitate sRNA–RNA interactions, like the Hfq and ProQ proteins (reviewed in (Updegrove *et al*., 2016, Santiago-Frangos & Woodson, 2018, Olejniczak & Storz, 2017)) or primarily bind mRNAs, such as CsrA (reviewed in (Romeo & Babitzke, 2018), with new classes in bacteria emerging, such as KH domain proteins (reviewed in (Olejniczak *et al*., 2021)).

A co-IP experiment is typically used to characterize RNA-binding proteins and identify bound RNAs. None of our co-IP experiments showed that BB0268 (FlgV) binds RNA, despite strong enrichment of both 3XFLAG-tagged and native FlgV protein in independent experiments. We analyzed the conservation of Hfq in bacteria and found no Hfq homologs in *B. burgdorferi*. Additionally, we were unable to reproduce previous results (Lybecker *et al*., 2010) that claimed *flgV* complements *E. coli* Δ*hfq* phenotypes. We also assayed the effect of expressing *E. coli hfq* in Δ*flgV B. burgdorferi* and found no complementation of the *flgV*-dependent defects in growth (Figure S4D and S4E). Caution generally should be taken when interpreting results from ectopic gene expression in disparate bacteria. Another study misannotated an RNA binding protein in *Pseudomonas aeruginosa* because the purified protein samples from *E. coli* extracts were contaminated with *E. coli* Hfq (Milojevic *et al*., 2013). Stress responses or other indirect effects also may be invoked from the expression of a foreign protein in a bacterium.

The effect of *flgV* on the RpoS regulator was examined both in *E. coli* and *B. burgdorferi*. Across bacteria, the activation of RpoS is complex, as it can involve multiple stress signals and is controlled at the level of transcription, translation and RNA/protein stability (reviewed in (Gottesman, 2019)). In *E. coli*, we observed some minor effects on RpoS levels when *flgV* was expressed; however, these outcomes could be indirect. In *B. burgdorferi*, RpoS is a cornerstone for responding to environmental changes during the enzootic cycle, and therefore, its regulation must be tightly controlled (Wachter *et al*., 2023). It was previously proposed that FlgV post-transcriptionally regulates *rpoS* because it was reported that deletion of *flgV* led to higher levels of *rpoS* mRNA, but lower levels of RpoS protein and lower levels the RpoS-regulated *ospC* mRNA (Lybecker *et al*., 2010). However, we found that when *rpoS* levels increased in Δ*flgV* samples there was a concomitant increase in RpoS protein, *ospC* mRNA and OspC protein (Figure S7A and S7B), opposite to what was reported previously (Lybecker *et al*., 2010). There were some variations in the *flgV-rpoS* effect between experiments (Figure S6B), which could be due to the precise state of the collected cells. Even across four WT *B. burgdorferi* samples, which all had similar cell densities by darkfield microscopy enumeration, there were some variations in *rpoS* levels (Figure S7B). None of our observations support a hypothesis (Lybecker *et al*., 2010) that FlgV post-transcriptionally regulates RpoS.

Given that *B. burgdorferi* has a limited number of characterized transcription factors (Subramanian *et al*., 2000), RNA-based regulation is an attractive mechanism for fine-tuning gene expression during the transmission and survival between tick and mammalian environments. Thus, future work is warranted to identify and properly characterize *bona fide* RNA chaperones in *B. burgdorferi*.

### FlgV is a broadly conserved flagellar protein

In Spirochaetota, endoflagella are critical for cell shape, motility and survival. For *B. burgdorferi*, spirochetes in the *Ixodes* tick midgut must sense a blood meal and migrate into a mammalian host and, likewise, *B. burgdorferi* inhabiting a mammal sense a feeding tick and migrate into that tick. Thus, to effectively maintain the enzootic cycle, the *B. burgdorferi* gene products which assemble, propel, and break flagellar motors must be tightly controlled. The regulatory mechanisms that control *B. burgdorferi* flagellar assembly and function are largely unknown and, intriguingly, the transcription of *B. burgdorferi* flagellar biosynthesis genes has been reported to be constitutive. We showed that *B. burgdorferi flgV* is transcribed as part of complex transcriptional units with surrounding genes in the “*flgB* superoperon”. The numerous and overlapping RNA transcripts in this region may have confounded the interpretations of experiments using reverse transcriptase-qPCR across ORF junctions (Zhang *et al*., 2020). Complex transcription, mRNA processing, and the presence of sRNAs across this operon may serve regulatory functions for the expression of specific genes, and thus control of motility, during the enzootic cycle.

There is an inherent link between the cell-cycle and flagella integrity for spirochetes (Lin *et al*., 2015, Motaleb *et al*., 2000), and we observed that when *flgV* is missing, spirochetes have defects in cell division and motility. Flagella-mediated motility is likely required to efficiently separate dividing spirochetes. We detected an abundance of conjoined spirochetes, and the presence of septa, for logarithmically growing Δ*flgV B. burgdorferi* in culture. In stationary phase, spirochetes lacking *flgV* were the same length as WT spirochetes. This suggests that in the absence of *flgV*, cells have prolonged cell division when rapidly growing, although ultimately division can take place. Cell elongation was also observed with *flgV* overexpression in logarithmic-phase spirochetes. Cell division genes (*ftsZ* (*bb0299*), *ftsA* (*bb0300*), *ftsW* (*bb0302*)) are encoded upstream of the “*flgB* superoperon”. Cross-talk between flagellar assembly and cell division also has been observed in other bacteria. Cell division-site placement in *Campylobacter jejuni* is influenced by *flhG*, the gene encoding a regulator of flagella number, (Balaban & Hendrixson, 2011). As in *B. burgdorferi*, *C. jejuni flhG* is encoded directly upstream of the *C. jejuni flgV* homolog. In *Caulobacter crescentus*, cell division is regulated in tandem with flagellar assembly (Siwach *et al*., 2021). Further work is warranted to explore the connection between cell division proteins and motility in *B. burgdorferi*.

We found *flgV* contributes to *B. burgdorferi* flagellar assembly. Δ*flgV* spirochetes have shorter and fewer flagellar filaments, compared to WT spirochetes. Yet strikingly, the number of motors is not impacted in Δ*flgV* spirochetes, nor is the overall structure of the flagellar motor. Only four other *B. burgdorferi* mutants have demonstrated such a flagellar filamentation defect. Deletions of *fliH* or *fliI*, encoding components of the flagellin export apparatus, result in short flagellar filaments, but do not prevent the formation of flagellar motors and filaments (Lin *et al*., 2015). Deletion of *fliD*, encoding the flagellar cap protein, or deletion of *flaB*, encoding the major flagellin protein, abolishes flagellar filamentation and results in short, disoriented flagellar hooks (Zhang *et al*., 2019).

Our study used cryo-ET to document that FlgV is an inner membrane protein, incorporated between the MS- and C-rings of the flagellar motor and sets the stage for further exploration of FlgV function and the control of motility. In general, once the flagellar basal body is assembled, flagellin is secreted to build the filament (Jones & Macnab, 1990, Hughes *et al*., 1993). Therefore, flagellin homeostasis and export of cytosolic flagellin into the basal body generally is co-regulated (Mukherjee *et al*., 2011, Oshiro *et al*., 2019). Typically, in model Enterobacteriaceae and Bacillaceae, σ^28^ controls the transcription of genes encoding the flagellar filament. *Borreliae* lack σ^28^. In this study, we showed that FlgV, which is strongly co-conserved with σ^28^ across other Spirochaetes and Firmicutes and other phyla, is another protein that modulates flagellar filamentation.

The stoichiometry of flagellar proteins is likely crucial for proper flagellar assembly. Yet, surprisingly, we observed no noticeable change to the motor structure when FlgV was overproduced, despite severe impacts to flagellar filamentation. FlgV harbors a conspicuous, conserved intramembrane basic residue (in a “RS” signature in *Borrelia*) in the second transmembrane helix, hinting that the protein could act as an intramembrane flagellar sensor. We hypothesize that environmental signals may promote/inhibit FlgV activity, which might be to tether/bring factors to the flagellar apparatus, perhaps through protein–protein associations at the FlgV C-terminal peptide tail.

We observed three distinct conservation patterns of FlgV in bacteria. We defined these as Type-1 in the Spirochaetota, PVC clade, Firmicutes, Nitrospinae and several other less-studied bacterial phyla, Type-2 in the *Helicobacter-* and *Campylobacter-*like Epsilonproteobacteria, and Type-3 in the *Arcobacter*-like Epsilonproteobacteria. Notably, the Epsilonproteobacteria (Type-2 and Type-3) FlgV differ from the Type-1 FlgV in that they lack: (i) the characteristic arginine in the second TM helix; (ii) an acidic residue at the start of TM helix-2; and (iii) a conserved polar residue towards the C-terminus of TM helix-2. Keeping with these differences, the Type-2 and Type-3 FlgV did not retrieve Type-1 FlgV with strong statistical significance. Nevertheless, structural comparisons revealed that they adopted a similar structure and they showed comparable gene-neighborhood associations within the flagellar operons. Simultaneous to our work, another group showed FlgV in *Helicobacter pylori* also associates with flagellar motors and impacts motility (Botting *et al*., 2023). We did not identify a FlgV homolog in flagellated-Enterobacteriaceae, all of which harbor σ^28^. Yet, there might be other functional analogs of FlgV that were not found by our analysis. The mechanistic details of how FlgV impacts flagellar filamentation and the differences across bacterial species are exciting directions for future work.

While endoflagella define spirochetes, there are notable differences (reviewed in (San Martin *et al*., 2023)). In Spirochaetes, *Borrelia*, which rely on a tick vector for survival, uniquely lack σ^28^. Constitutive expression of flagellar genes may be required for a spirochete to quickly respond to a sporadic bloodmeal and migrate into a mammalian host. Other forms of regulation including the modulation of flagellar filamentation, perhaps through *flgV*, is likely critical for energy conservation and increased motility during a bloodmeal. A limited number of flagella are required for maintaining cell shape in *Borrelia*, but there may be times during the enzootic cycle such as in an unfed tick, where some flagellar motors lack a filament. Future experiments to examine FlgV and other proteins encoded by the heterogeneous flagellar superoperons likely will reveal new insights into the regulation of these important nanomachines, which impact motility, morphology, cell division and infectivity across the bacterial superkingdom.

### Importance of *B. burgdorferi* motility during infection

Pathogens must overcome numerous physical conditions, nutrient limitations and immune defenses to survive inside a host. *B. burgdorferi*, with a small, segmented genome and no known virulence factors, heavily relies on motility to move from host-to-host, scavenge for nutrients and avoid immune factors. We undertook a unique opportunity to investigate the importance of motility during infection by testing Δ*flgV* and ++*flgV* spirochete survival during tick and mammalian infection. On average, we found WT spirochetes to harbor 9 filaments, Δ*flgV* spirochetes to harbor 7 filaments, and ++*flgV* spirochete to harbor 3 filaments. Both Δ*flgV* and ++*flgV B. burgdorferi* survived and replicated in ticks at the same levels as WT *B. burgdorferi*. This suggests that motility is not critical for *B. burgdorferi* inside ticks. A previous study observed that *B. burgdorferi* lacking flagellar filaments entirely (Δ*flaB*) can survive, but at a reduced burden, in ticks (Sultan *et al*., 2013). Yet, Δ*flgV* and ++*flgV B. burgdorferi* were attenuated in their ability to infect mice.

Successful mammalian infection of *B. burgdorferi* begins with a localized skin infection of spirochetes at the tick bite site, followed by dissemination and long-term colonization of distal tissues. We observed a reduction in Δ*flgV B. burgdorferi* expansion in the skin inoculation site, during the first week of infection by needle inoculation. This was followed by attenuation of hematogenous spread and a delay in Δ*flgV* spirochetes colonizing distal tissues. When *flgV* was overexpressed spirochetes did not survive mouse infection and likely were eliminated at the skin inoculation site before dissemination. The infection phenotypes associated with altered *flgV* levels may be explained by the innate immune response effectively eliminating less-motile spirochetes, the inability of *flgV-*mutant spirochetes to migrate through physical barriers (connective tissue, bloodstream wall, etc.), or both. Interestingly, the Δ*flgV B. burgdorferi* that do survive the initial steps in infection and have a slower kinetics of tissue colonization compared to WT spirochetes, ultimately colonize heart and joint tissue at a similar level to WT spirochetes. Yet, we observed a strong attenuation of *flgV-*mutant spirochetes for colonization of ear tissue. It was previously reported that Δ*bb0268* (*flgV*) *B. burgdorferi* are not infectious, as no Δ*bb0268* (*flgV*) spirochetes were detected from reisolation attempts using ear punches at 3 weeks or using ear punches, joints and bladders at 5 weeks post-intraperitoneal needle inoculation of spirochetes (Lybecker *et al*., 2010). However, infecting mice by intradermal needle injection is more physiologically similar to a natural *B. burgdorferi* infection by tick-bite, than intraperitoneal needle injection, which may account for differences between the previous infection study (Lybecker *et al*., 2010) and our data. Some studies have suggested that motility is absolutely required for *B. burgdorferi* infectivity (Lin *et al*., 2012, Sultan *et al*., 2013, Lin *et al*., 2015). However, our findings suggest that once spirochetes have disseminated to a tissue, motility may not be as critical for survival compared to initial skin-localized *B. burgdorferi* infection at the inoculation site.

In all, this work encompasses the first investigation of FlgV as a modulator of flagellar assembly in bacteria. We tested the consequences of defective motility during tick and mammalian infection, revealing even slight changes to the number/length of flagellar filaments have significant consequences for *B. burgdorferi* infectivity. Experiments to test when FlgV, and other proteins that modulate motility, are active during tick-mammalian infection, will be critical to define the hierarchical, flagellar regulatory network in *B. burgdorferi*. As we gain insights into how spirochetes traverse, replicate and divide inside a host, we will better understand the biology of Lyme disease.

## MATERIALS AND METHODS

### Bacterial strains and plasmids

Derivatives of *E. coli* K12 MG1655 (WT) or *B. burgdorferi* B31 A3 (WT; Adams lab strain number PA003, which lacks cp9) were used for all experimental studies. The genetically tractable and infectious *B. burgdorferi* B31 A3-68Δ*bbe02* strain (WT; Adams lab strain number PA001, derivative of PA003, which lacks cp9, lp56 and gene *bbe02* on lp25 (Rego *et al*., 2011)), was used for chromosomal mutants and plasmid transformations, as indicated. All strains, plasmids, oligonucleotides, and synthesized DNA sequences used are listed in Table S1. Chromosomal mutants were verified by PCR and sequencing and/or whole genome sequencing. Engineered plasmid inserts were verified by sequencing. All *B. burgdorferi* transformants were verified by PCR to contain the endogenous plasmid content of the parent strain, using a panel of primers (Elias *et al*., 2002).

All *B. burgdorferi* chromosomal mutations were performed by allelic exchange. The *B. burgdorferi bb0268*(*flgV*)-3XFLAG construct, harboring a spectinomycin/streptomycin (*flaBp-aadA*) resistant cassette, was PCR amplified using primers MJ1049 + MJ1050, PA102 + PA103, MJ1051 + MJ1052 with an overlap extension strategy (Ellis *et al*., 2014). The final PCR product was ligated with linear pCR-Blunt using a Zero Blunt PCR cloning kit (Life technologies) in DH5ɑ *E. coli*. Twenty micrograms of pCR-Blunt-*flgV*-3XFLAG were transformed into *B. burgdorferi* B31 WT (PA003), resulting in strain PA007. Two independent *B. burgdorferi* Δ*flgV*(*bb0268*) strains were created for this study. Allelic exchange constructs were generated using primers PA337 + PA352, PA102 + PA103, PA353 + PA340 and the final PCR-generated linear DNA was either ligated into pCR-Blunt (as above) or directly transformed into WT *B. burgdorferi* (PA001), resulting in Δ*flgV* strains PA011 and PA251, respectively. The *gfp-flgV B. burgdorferi* strain was made by gene synthesis of the allelic exchange construct (*flgV*-ORF fused to *gfp*, *flaBp-aadA*, and 500 bp of the surrounding genomic region; Table S1) and cloned into pUC57 (GenScript). A previously published *gfp* sequence (Takacs *et al*., 2018), codon-optimized for *B. burgdorferi* was used. PCR from this plasmid (primers PA667+PA672) was performed and 20 μg of the PCR-product was directly transformed into WT *B. burgdorferi* (PA001), resulting in strain PA402. The *B. burgdorferi* Δ*rpoS* strain, harboring a gentamycin (*flgBp-aacC1*) resistant cassette, was PCR amplified using primers PA411 + PA412, PA415 + PA416, PA413 + PA414 and the final PCR product was directly transformed into WT *B. burgdorferi* (PA001), resulting in strain PA235.

All engineered plasmids utilized the *B. burgdorferi* shuttle vector, pBSV2G (Elias *et al*., 2003). To generate an *E. coli hfq* expression vector in *B. burgdorferi*, the *E. coli* Hfq ORF was codon-optimized for *B. burgdorferi* with the OptimumGene^TM^ algorithm and synthesized (Table S1). The codon-optimized gene was PCR amplified with primers PA154 + PA155. These PCR products were subsequently cut with *Nde*I and *BamH*I restriction enzymes and ligated into a pBSV2G derivate that harbors a constitutively active *flaB* promoter (Jain *et al*., 2015). This resulted in p_con_+*E. coli hfq*. To generate a *flgV* overexpression strain (++*flgV*), the native *B. burgdorferi flgV* ORF was PCR amplified with primers PA156 + PA157, using *B. burgdorferi* A3 genomic DNA as template. These PCR products were subsequently cut with *Nde*I and *BamH*I restriction enzymes and ligated into a pBSV2G derivate that harbors a constitutively active *flaB* promoter (Jain *et al*., 2015). This resulted in p_con_++*flgV*. To create an IPTG inducible plasmid on pBSV2G (p_ind_), we inserted the *flaB promoter-lacI* and *pQE30* promoter from pJSB252 (Blevins *et al*., 2007) onto empty pBSV2G and replaced the *flaB* promoter on p_con_++*flgV* using NEBuilder HiFi DNA Assembly (New England Biolabs) to create p_ind_ and p_ind_+*flgV*, respectively. Briefly, high-fidelity PCR reactions (NEB Q5 DNA Polymerase) with pBSV2G and p_con_++*flgV* were performed using primers PA470 + PA467 and PA466 + PA467, respectively, and the resultant PCR product gel extracted using a NucleoSpin Gel and PCR Clean-up kit (Macherey-Nagel). *flaB* promoter*-lacI* and *pQE30* were Q5 PCR-amplified from pJSB252 using primers PA468 + PA471 (for p_ind_ template) and PA468 + PA469 (for p_con_++*flgV* template) and mixed with the corresponding linear, gel-purified vector at a 2:1 ratio in a 20 μl NEBuilder reaction at 50°C for 1 h. Ten microliters were subsequently transformed into NEB Turbo Competent *E. coli*.

### Growth conditions

*E. coli* strains were grown with shaking at 250 rpm at 37°C in LB rich medium supplemented with 10 μg/ml gentamycin when applicable. Overnight *E. coli* cultures were diluted to an OD_600_ of 0.05 and grown to the indicated time point.

*B. burgdorferi* were cultivated in liquid Barbour-Stoenner-Kelly (BSK) II medium supplemented with gelatin and 6% rabbit serum (Barbour, 1984) and plated in solid BSK medium, as previously described (Rosa & Hogan, 1992). *B. burgdorferi* cultures were grown at 35°C with 2.5% CO_2_ in the presence of 50 μg/ml streptomycin or 40 μg/ml gentamycin when applicable. IPTG (100 μM) was added at the subculture for clones harboring p_ind_ derivates. *B. burgdorferi* starter cultures were subcultured only once from the freezer stock, by diluting spirochetes at 1×10^5^ cells/ml, for all experiments.

### Growth analysis

*E. coli* growth curves were performed in biological triplicate, by measuring the OD_600_ every 60 min for 780 min total, after a dilution of the overnight culture to an OD_600_ of 0.05. *B. burgdorferi* growth curves were performed in biological triplicate, by diluting spirochetes to 1×10^5^ cells/ml and monitoring cell growth at the indicated time points by darkfield microscopy enumeration.

### RNA-Protein coimmunoprecipitation (Co-IP) assay

Co-IP of Hfq, BB0268 (FlgV) and KhpB proteins were performed using the initial steps of the RIL-seq protocol (Melamed *et al*., 2018). Samples were crosslinked with 80,000 μJ/cm^2^ of UV irradiation, lysed with 0.1 mm glass beads, and co-IPs carried out using 3 μg monoclonal anti-FLAG M2 antibody (Sigma-Aldrich) or 30 μg polyclonal FlgV antibody or 30 μg polyclonal KhpB antibody and 20 μl of Pierce protein A/G magnetic beds. Polyclonal antibodies to KhpB were generated by immunizing rabbits with purified His-tagged recombinant KhpB protein, followed by affinity purification (GenScript). Aliquots of total lysate, bead supernatant, and IP beads were mixed with equal volume 2X Laemmli sample buffer (Bio-Rad) and subjected to immunoblot analysis as below. Aliquots of total lysate and bead supernatant were mixed with TRIzol (Thermo Fisher Scientific) and RNA extracted according to the standard protocol. RNA elution buffer (50 mM Na_3_PO_4_, 300 mM NaCl, 300 mM Imidazole, 0.1 U/μl recombinant RNase inhibitor) was added to the IP beads prior to addition of TRIzol and RNA extraction. RNA from all samples was resuspended in 12 μl of DEPC H_2_O and analyzed using an Agilent 4200 TapeStation System or northern blot analysis, as described below. In addition to the data included in this manuscript we varied the amount of *B. burgdorferi* cells, growth phase and UV intensity for cross-linking, but all trials resulted in no enrichment of RNA with BB0268.

### RNA isolation

*E. coli* cells corresponding to the equivalent of 10 OD_600_ or ∼45 ml of a *B. burgdorferi* culture were collected by centrifugation, washed once with 1X PBS (1.54 M NaCl, 10.6 mM KH_2_PO_4_, 56.0 mM Na_2_HPO_4_, pH 7.4) and pellets snap frozen in liquid N_2_. RNA was isolated using TRIzol (Thermo Fisher Scientific) exactly as described previously (Melamed *et al*., 2020). RNA was resuspended in 20–50 µl DEPC H_2_O and quantified using a NanoDrop (Thermo Fisher Scientific).

### Northern blot analysis

Northern blots were performed using total RNA exactly as described previously (Melamed *et al*., 2020). For small RNAs, 5 μg of RNA were fractionated on 8% polyacrylamide urea gels containing 6 M urea (1:4 mix of Ureagel Complete to Ureagel-8 (National Diagnostics) with 0.08% ammonium persulfate) and transferred to a Zeta-Probe GT membrane (Bio-Rad). For longer RNAs, 10 μg of RNA were fractionated on a 2% NuSieve 3:1 agarose (Lonza), 1X MOPS, 2% formaldehyde gel and transferred to a Zeta-Probe GT membrane (Bio-Rad) via capillary action overnight. For both types of blots, the RNA was crosslinked to the membranes by UV irradiation. RiboRuler High Range and Low Range RNA ladders (Thermo Fisher Scientific) were marked by UV-shadowing. Membranes were blocked in ULTRAhyb-Oligo Hybridization Buffer (Ambion) and hybridized with 5′ ^32^P-end labeled oligonucleotides probes (listed in Table S1). After an overnight incubation, the membranes were rinsed with 2X SSC/0.1% SDS and 0.2X SSC/0.1% SDS prior to exposure on film. Blots were stripped by two 7-min incubations in boiling 0.2% SDS followed by two 7-min incubations in boiling water.

### Immunoblot analysis

Polyclonal antibodies to FlgV, FlaB, and RpoS were generated by immunizing rabbits with purified His-tagged recombinant FlgV protein (amino acids 58–159), recombinant FlaB, or recombinant RpoS protein followed by affinity purification (GenScript). Immunoblot analyses using *B. burgdorferi* lysates and probing for FlgV (Figure 3A), FlaB (Figure S7D) and RpoS (Figure S7E) confirmed the specificity of the antibodies generated in this study. *E. coli* immunoblot analyses were performed as described previously (Zhang *et al*., 2002), with minor changes listed below. *B. burgdorferi* total protein lysates were collected by centrifugation and cell pellets were washed twice with 1X HN (50 mM Hepes, 50 mM NaCl, pH 7.5), before boiling with Laemmli Sample Buffer + β-mercaptoethanol (Bio-Rad). Samples were separated on a Mini-PROTEAN TGX 4%–20% Tris-Glycine gel (Bio-Rad) and transferred to a nitrocellulose membrane (Thermo Fisher Scientific). Membranes were blocked in 1X TBST containing 5% milk and probed with a 1:2,000 dilution of monoclonal α-FLAG-HRP (Sigma-Aldrich), 1:5,000 dilution of *E. coli* α-RpoS (gift from Sue Wickner, NIH), 1:5,000 dilution of α-Hfq (Zhang *et al*., 2002), 1:500 dilution of α-FlgV (this study), 1:200 dilution of α-FlaB (Barbour *et al*., 1986) or 1:20,000 dilution of α-FlaB (this study), 1:1,000 dilution of α-FlgG (Zhang *et al*., 2020, Zhao *et al*., 2013), 1:2,000 dilution of α-FliF (Zhang *et al*., 2020), 1:500 dilution of α-RpoS (this study), 1:1,000 dilution of α-OspC (Rockland), or 1:2,000 dilution of α-SodA (Esteve-Gassent *et al*., 2009). Secondary peroxidase labeled α-mouse (GE Healthcare), α-rabbit (GE Healthcare) or α-rat (Invitrogen) HRP-conjugated antibodies were subsequently used at a 1:10,000 dilution, when required. Membranes were developed with either SuperSignal West Pico PLUS or West Femto Maximum Sensitivity Chemiluminescent Substrate (Thermo Fisher Scientific) on a Bio-Rad ChemiDoc MP Imaging System. Membranes were stripped for 15 min with Restore Immunoblot Blot Stripping Buffer (Thermo Fisher Scientific).

### Protein sequence and structure analysis

Sequence-profile searches were performed with the PSI-BLAST program against the NCBI non-redundant (nr) database clustered down to 50% sequence identity using the MMSEQS program with a profile-inclusion threshold was set at an e-value of 0.01. Iterative HMM searches were performed using the JACKHMMER program from the HMMER3 package. Profile-profile searches against PDB, Pfam and an in-house collection of profiles maintained by the Aravind lab were performed using the HHpred program. The FAMSA and MAFFT programs were used to construct multiple sequence alignments (MSAs). TM segments and signal peptides were predicted using a deep neural network as implemented in SignalP 6 and HMM-based TMHMM (Krogh *et al*., 2001) and Phobius programs.

Secondary structures were inferred using the JPred3 program with MSAs as inputs. PDB coordinates of structures were retrieved from the Protein Data Bank. Structural models were generated using the RoseTTAfold and Alphafold2 programs (Jumper *et al*., 2021) utilizing the NIH Biowulf cluster to run the GPU-dependent steps. Multiple alignments of related sequences (>30% similarity) were used to initiate HHpred searches for the step of identifying templates to be used by the neural networks deployed by these programs.

Clustering of protein sequences was done using the MMSEQS program by empirically adjusting the length of aligned regions and bit-score densities. Phylogenetic analysis was performed using the maximum-likelihood method with the LG or JTT models with the FastTree program. The same program was used to generate distance matrices used in separating groups based on differential evolutionary rates. A custom PERL script was used to extract gene neighborhoods from genomes retrieved from the NCBI Genome database. These were then clustered using MMSEQS and filtered using a neighborhood distance cutoffs and phyletic patterns to identify conserved gene neighborhoods.

The output from the MSAs and phylogenetic analyses are published as metadata (temporally available at FigShare: https://figshare.com/s/7ca8d928d037bb358e02).

### *B. burgdorferi* cell length analysis

*B. burgdorferi* were collected by centrifuged at 2000 x g for 10 min at 25°C and washed twice in sterile 1X PBS. Cell pellets were resuspended in 1X PBS and 3 μl were immobilized on a glass slide (FisherScientific) with 18×18-1.5 cover glass (FisherScientific). *B. burgdorferi* were imaged using Zeiss Axiolab 5 microscope with a 40X/0.65 NA objective. Images were captured using an Axiocam 305 color camera (Zeiss) and cells were measured using the curve (spline) tool with ZEN 3.4 (blue edition) software.

### *B. burgdorferi* motility plate assay

Motility plate assay was adapted from (Zhang & Li, 2018). On the day of the assay, a solution of 0.625% agarose was autoclaved and combined with *B. burgdorferi* plating medium for a final agarose concentration of 0.35%. 36 ml of the combined medium and agarose was added to a petri dish and left to dry in a biosafety cabinet for 1 h. A 10 ml subculture of *B. burgdorferi* was grown to logarithmic phase. Cells were centrifuged at 2000 x g for 20 min, resuspended in BSKII to a final density of ∼5×10^5^ cells/µl, and 2 µl of cell suspension were inoculated just under the agarose surface. The plates were left uncovered in a biosafety cabinet for 30 min and then incubated at 35°C with 2.5% CO_2_ for 9 days. The diameters of the motility rings were measured using a ruler, measuring two perpendicular diameters for each ring and calculating the mean for each sample. Measurements were made to the nearest 1 mm.

### FlgV localization

*B. burgdorferi* protein lysates were separated into soluble and membrane fractions, as performed in (Kuhn *et al*., 2021). Briefly, spirochetes were grown to log phase (∼2.2×10^7^ cells/ml) in BSKII medium, spun at 3210 x g for 10 min, washed twice with cold 1X HN buffer, and resuspended in 1 ml 1X HN with 100 μl Halt Protease Inhibitor Cocktail (Thermo Fisher Scientific). Spirochetes were lysed by sonication and spun at 125,000 x g to collect the soluble and membrane components of the lysate.

### Fluorescence microscopy

*B. burgdorferi* were centrifuged at 2000 x g for 10 min at 25°C and resuspended in sterile 1X PBS at a final density of ∼1×10^5^ cells/µl. 3 µl of cells were immobilized on a 35 mm glass-bottom dish (MatTek) with a 1 cm disc of 0.77% agarose. *B. burgdorferi* were imaged using Zeiss LSM800 confocal microscope with a 63X/1.4 NA oil objective. Green fluorescence and bright-field images were captured for each field-of-view. Images were processed with software Fiji ImageJ (version 2.1.0/1.53c).

For demography analysis, images were analyzed with the software Fiji ImageJ2 (version 2.3.0/1.53q). The fluorescence intensity was plotted free-hand along each cell’s length using the Plot Profile tool on the fluorescent channel, yielding a data frame for each cell. A total of 300 cells were analyzed for each growth condition. Using R (version 4.2.1) and RStudio (version 2022.07.0, build 548), the data frames were organized and depicted in a demograph using the tidyverse (Wickham *et al*., 2019), dplyr (Wickham *et al*., 2023), and ggplot2 packages (Wickham, 2016).

### Cryo-ET sample preparation, data collection, and reconstruction

*B. burgdorferi* were grown to a density of: 3.5 x 10^7^ cells/ml for WT/p (PA023), 1.05 x 10^7^ cells/ml for WT/p_con_++*flgV* (PA267), 4.5 x 10^6^ cells/ml for WT/p_ind_ (PA273), 7.5 x 10^5^ cells/ml for Δ*flgV*/p_ind_ (PA310), 6.0 x 10^6^ cells/ml for Δ*flgV*/p_ind_+*flgV* (PA312), and 2.0 x 10^7^ cells/ml WT *flgV-gfp* (PA402). Cultures were shipped overnight to the Microbial Sciences Institute at Yale University, where they were immediately processed. Cultures were centrifuged at 5,000 rpm and washed three times with 1X PBS. 10 µl of resuspended *B. burgdorferi* (normalized to an OD_600_ = 1.0) were mixed with 20 µl 10-nm colloidal gold particles. The mixture was deposited onto freshly glow-discharged grids (R2/1 Quantifoil) for 1 min, blotted with filter paper for 3 seconds, and frozen in liquid ethane using a gravity-driven homemade plunger apparatus, as described previously (Liu *et al*., 2009).

Bacterial samples were imaged with a Titan Krios microscope (Thermo Fisher Scientific) equipped with a field emission gun, an energy filter, and a K3 direct-detection device (Gatan). An energy filter with a slit width of 20 eV was used for data acquisition at an -6 μm defocus.

To determine the flagellar motor structures, we only collected tilt series from the cell poles at magnification of x42,000, resulting in a pixel size of 2.148 Å. For each tilt series, we collected 33 image stacks at a range of tilt angles of between +48° and −48° (3° step size) using a bidirectional scheme with a cumulative dose of 60e^-^/Å^2^. All the title series were aligned and reconstructed by IMOD (Kremer *et al*., 1996, Mastronarde & Held, 2017). In total, we generated 120 tomograms from WT/p (PA023), 64 from Δ*flgV*/p_ind_ (PA310), 41 from Δ*flgV*/p_ind_+*flgV* (PA312), 190 from WT *flgV-gfp* (PA402), and 65 from WT/p_con_++*flgV* (PA267) *B. burgdorferi*.

To visualize flagella and cell poles for over 2 μm length, multiple tilt series were collected along the spirochetes by SerialEM (Mastronarde, 2005). For each tilt series, we collected 33 image stacks at a range of tilt angles of between +48° and −48° (3° step size) using a bidirectional scheme with a cumulative dose of 80e^-^/Å^2^ at magnification of x19,000, resulting in a pixel size of 4.556 Å at the specimen level. All the title series were aligned and reconstructed by IMOD (Kremer *et al*., 1996, Mastronarde & Held, 2017). In total, we generated 21 tomograms from WT/p_ind_ (PA273), 21 from Δ*flgV*/p_ind_ (PA310), 18 from Δ*flgV*/p_ind_+*flgV* (PA312), 18 tomograms from WT/p (PA023), and 18 from WT/p_con_++*flgV* (PA267) *B. burgdorferi*. Flagellar motors and flagellar filaments were manually tallied for each tomogram.

### Cryo-ET data visualization

Representative reconstructions from *B. burgdorferi* strains PA023, PA310, PA312 and PA267 and were visualized in IMOD (Kremer *et al*., 1996). Periplasmic flagella, the motors, the outer membranes, and inner membranes were manually segmented in IMOD.

### Subtomogram averaging

To determine flagellar motor structures by subtomogram averaging, we visually identified the motors from the tomograms. In total, we found 907 motors from PA023, 562 from PA310, 272 from PA312, 1462 from PA402, and 485 from PA267. I3 package (Winkler *et al*., 2009) was used to determine *in situ* structures as described previously (Liu *et al*., 2009).

### Ethics statement

The National Institutes of Health and the University of Central Florida are accredited by the International Association for Assessment and Accreditation of Laboratory Animal Care. All animals were cared for in compliance with the Guide for the Care and Use of Laboratory Animals. Protocols for all animal experiments were prepared according to the guidelines of the National Institutes of Health, reviewed and approved by the *Eunice Kennedy Shriver* National Institute of Child Health and Human Development Institutional Animal Care and Use Committee or the UCF Institutional Animal Care and Use Committee.

### Infection of *Ixodes* larvae by immersion and tick feeding

Approximately 2-month-old *Ixodes scapularis* naïve larval ticks (Oklahoma State University, Department of Entomology and Plant Pathology) were infected by immersion (Policastro & Schwan, 2003). Larval ticks were dehydrated by exposure to saturated ammonium sulfate for ∼88 h. *B. burgdorferi* strains were grown to logarithmic phase (2×10^7^ – 5.3×10^7^ cells/ml) and diluted to 2×10^7^ cells/ml in BSKII. 500 μl of spirochetes were incubated with dehydrated ticks at 35°C for 1.5 hours and washed twice with sterile 1X PBS. The inoculum cultures were verified to contain the expected endogenous plasmids, using a panel of primers (Elias *et al*., 2002). Individuals from each inoculum clone were analyzed for the presence of virulence plasmids lp25, lp28-1 and lp36 (Jewett *et al*., 2009), each of which was confirmed to be present in 80– 100% of the population.

Twelve days following the artificial infection, groups of 10 unfed larvae were surface sterilized and plated in solid BSKII containing RPA cocktail (60 μM rifampicin, 110 μM phosphomycin, 2.7 μM amphotericin B) and 40 μg/ml gentamycin to determine *B. burgdorferi* CFUs/10 larvae. Two days following the artificial infection, groups of ∼129 artificially infected larvae, on average, were fed to repletion on groups of 3 naïve, 6–8-week-old, C3H/HeN mice (Envigo). Mice were assayed for *B. burgdorferi* infectivity by reisolation in BSKII, as described in the section below. Ticks fed for ∼1 week before naturally falling off. Approximately 1 week following the tick collection, individual fed larvae were analyzed for *B. burgdorferi* burden by plating, as described above. A subset of fed larvae were allowed to molt into nymphs, and ∼5 weeks following, individual unfed nymphs were analyzed for *B. burgdorferi* burden by plating, as described above. Groups of ∼5 unfed nymphs, on average, were fed on groups of 3 naïve, 6–8-week-old, C3H/HeN mice (Envigo) to repletion. Ticks fed for ∼1 week before naturally falling off. Approximately 1 week following the tick collection, individual fed nymphs were analyzed for *B. burgdorferi* burden by plating, as above.

### Mouse infection by needle inoculation

All *B. burgdorferi* strains were grown to stationary phase (∼2-3×10^8^ cells/ml). Groups of 6 C3H/HeN mice (Envigo) each were needle inoculated with 1×10^4^ *B. burgdorferi* intradermally. Inoculum densities were confirmed by plating for individuals in solid medium. All inoculum cultures were verified by PCR to contain the expected endogenous plasmids (Elias *et al*., 2002, Jewett *et al*., 2007). Individuals from each inoculum clone were analyzed for the presence of virulence plasmids lp25, lp28-1 and lp36 (Jewett *et al*., 2009), each of which was confirmed to be present in 90–100% of the population.

### Determination of *B. burgdorferi* density in blood

On day 6 post-inoculation approximately 50 µl of blood was collected from each mouse into K2 EDTA MiniCollect blood collection system tubes (Greiner). Each 50 µl whole blood sample was added to 1 ml BSKII then plated in solid BSK-agarose medium plus RPA cocktail (60 µM rifampicin, 110 µM phosphomycin, and 2.7 µM amphotericin B) and plates incubated at 35°C under 2.5% CO_2_ for 7–14 days. Colony-forming units (CFU) were enumerated and *B. burgdorferi*/ml of blood was calculated as the CFU/volume of blood analyzed multiplied by 100 for each sample (Aranjuez *et al*., 2019).

### Spirochete reisolation and quantification of *B. burgdorferi* densities in tissues

Mouse infection was determined by cutting each mouse tissue in half and assaying spirochete reisolation and quantitative PCR for *B. burgdorferi* loads for skin inoculation site, ear, heart and joint samples (exactly as described previously (Jewett *et al*., 2007, Aranjuez *et al*., 2019)). TaqMan probe and primers (IDT DNA) specific to the *B. burgdorferi flaB* gene (primers: MJ1137, MJ1138, MJ1139) and mouse *nid* gene (primers: MJ1140, MJ1141, MJ1142) (Table S1) were used to measure their respective copy numbers by qPCR and a standard curve approach, using 100 ng of DNA extracted from mouse tissues as the template.

## ACKNOWLEDGMENTS

Thank you to the late C. Savage, a tenacious and inspiration Borreliologist and colleague, for pointing out that *bb0268* is encoded near flagellar genes. Thank you to J. Botting for indicating to us that BB0268 is a homolog to FlgV in *H. pylori*. Thank you to S. Gottesman, S. Melamed and members of the Adams and Storz lab for helpful comments on the manuscript. Thank you to A. Zhong for assistance with the AlphaFold2 analysis and J. Silberman for assistance with the FlaB and RpoS immunoblots. We also thank T. Li and J. Iben for whole genome sequencing and analysis. Thank you to J. Seshu for providing the ɑ-SodA antibodies, C. Li for the ɑ-FlgG and ɑ-FliF antibodies and M. Motaleb for the Δ*flaB* and Δ*flaB+flaB B. burgdorferi*. This work utilized the computational resources of the NIH HPC Biowulf cluster (http://hpc.nih.gov).

## Author contributions

P.P.A., G.S., and M.W.J conceived the project. P.P.A., G.S., M.W.J., J.L, and L.A. funded the project. P.P.A., G.S., M.W.J., J.L, and L.A. designed the experiments. M.Z.C., D.S., P.P.A., H.Y., A.G.L., Y.Y.C, and L.A. performed the experiments. P.P.A., G.S., M.W.J., L.A., and J.L. wrote the manuscript, with contributions from all authors. All authors critiqued and edited the final manuscript.

## MULTIMEDIA FILES

**Supplementary Video 1.** Three-dimensional reconstruction from a WT/p (PA023) *B. burgdorferi*.

**Supplementary Video 2.** Three-dimensional reconstruction from a Δ*flgV*/p_ind_ (PA310) *B. burgdorferi*.

**Supplementary Video 3.** Three-dimensional reconstruction from a Δ*flgV*/p_ind_+*flgV* (PA312) *B. burgdorferi*.

**Supplementary Video 4.** Three-dimensional reconstruction from a WT/p_con_++*flgV* (PA267) *B. burgdorferi*.

