## Supplemental Information for "Broadly conserved FlgV controls flagellar assembly and *Borrelia burgdorferi* dissemination in mice"

### TABLE OF CONTENTS

**Supp. Figure S1. BB0268-3XFLAG and native BB0268 does not bind RNA** (for Figure 1).

**Supp. Figure S2. The conserved *hfq* and *miaA* gene neighborhoods** (for Figure 1).

**Supp. Figure S3. AlphaFold secondary structure predictions for *E. coli* Hfq and *B. burgdorferi* BB0268 (FlgV)** (for Figure 1).

**Supp. Figure S4. BB0268 (FlgV) does not complement an *E. coli* Δ*hfq* strain and *E. coli* Hfq does not complement a *B. burgdorferi* Δ*bb0268* (*flgV*) strain** (for Figures 1 and 3).

**Supp. Figure S5. *bb0268* (*flgV*) is co-transcribed with flagellar genes** (for Figure 2).

**Supp. Figure S6. FlgV is a conserved, membrane-associated protein** (for Figures 2 and 4)

**Supp. Figure S7. FlgV levels impact FlaB protein, but not *flaB* mRNA levels** (for Figure 5).

### SUPPLEMENTARY FIGURES

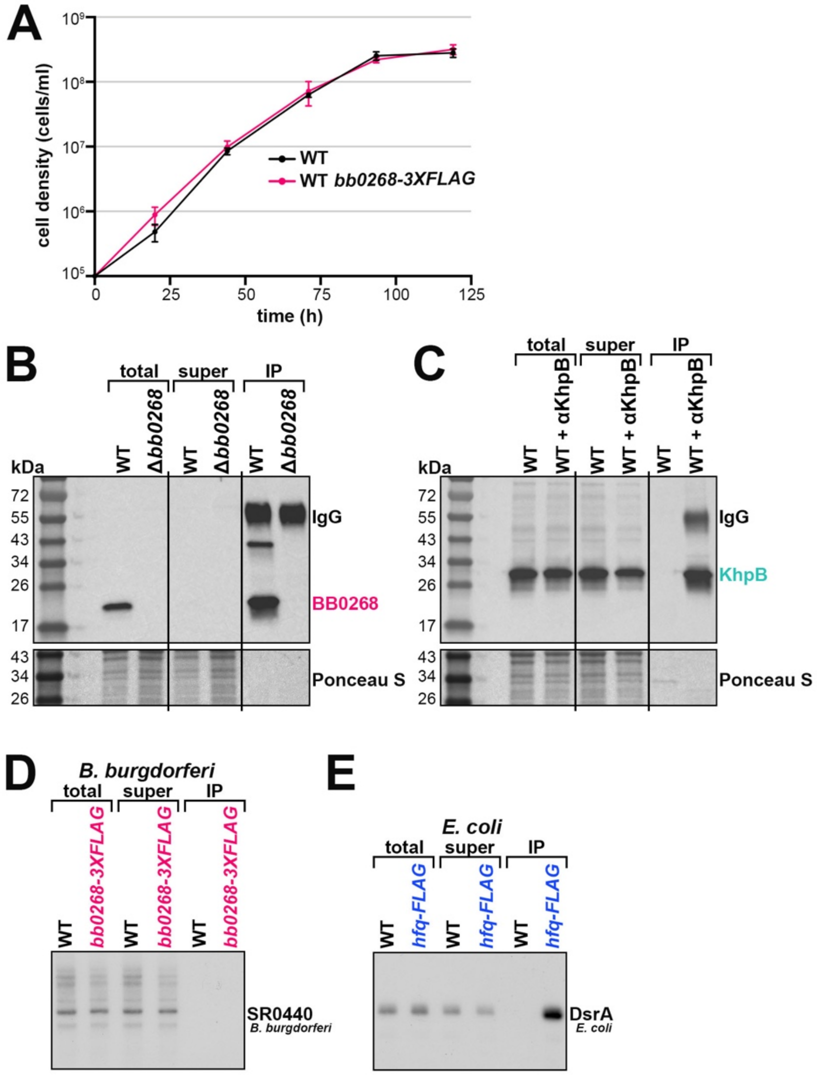


**Supp. Figure S1. BB0268-3XFLAG and native BB0268 does not bind RNA** (for Figure 1).

(**A**) Growth curve of *bb028-3XFLAG* (PA007) and WT (PA003) *B. burgdorferi*. Cell growth was monitored by dark field microscopy enumeration at the indicated time points, after dilution of the starter culture to $1x10^5$ cells/ml. Each data point (circles) represents the mean of three biological replicates with the standard deviation.

(**B**) Immunoblot analysis of immunoprecipitated WT *B. burgdorferi* (PA001) or Δ*bb0268 B. burgdorferi* (PA251) samples. *B. burgdorferi* cells were grown to $\sim 6.3x10^7$ cells/ml and then exposed to UV to crosslink any RNAs associated with the proteins. After cell lysis, the tagged proteins were immunoprecipitated (IP) and RNA was isolated from the IP samples. Equal volumes of cell lysis (total), supernatant of the α-BB0268-beads during washing, and elution from the α-BB0268-beads (IP) were separated on a Tris-Glycine gel, transferred to a membrane, stained with Ponceau S, and probed using α-BB0268 antibodies. Size markers are indicated. The IgG label denotes the heavy chain of the BB0268 antibody.

(**C**) Immunoblot analysis of immunoprecipitated WT *B. burgdorferi* (PA001) using KhpB antibodies. *B. burgdorferi* cells were grown to ~$8.1x10^7$ cells/ml and co-IP performed, as in panel B. The immunoblot was probed using α-KhpB antibodies. Size markers are indicated. The IgG label denotes the heavy chain of the KhpB antibody.

**(D)** Northern analysis of BB0268-3XFLAG RNA samples. One microliter of each *B. burgdorferi* sample from Figure 1A was separated on an acrylamide gel, transferred to a membrane, and probed for *B. burgdorferi* DsrA.

(**E**) Northern analysis of *E. coli* Hfq-FLAG RNA samples from Figure 1A. Samples were analyzed as in panel D and probed for *E. coli* DsrA.

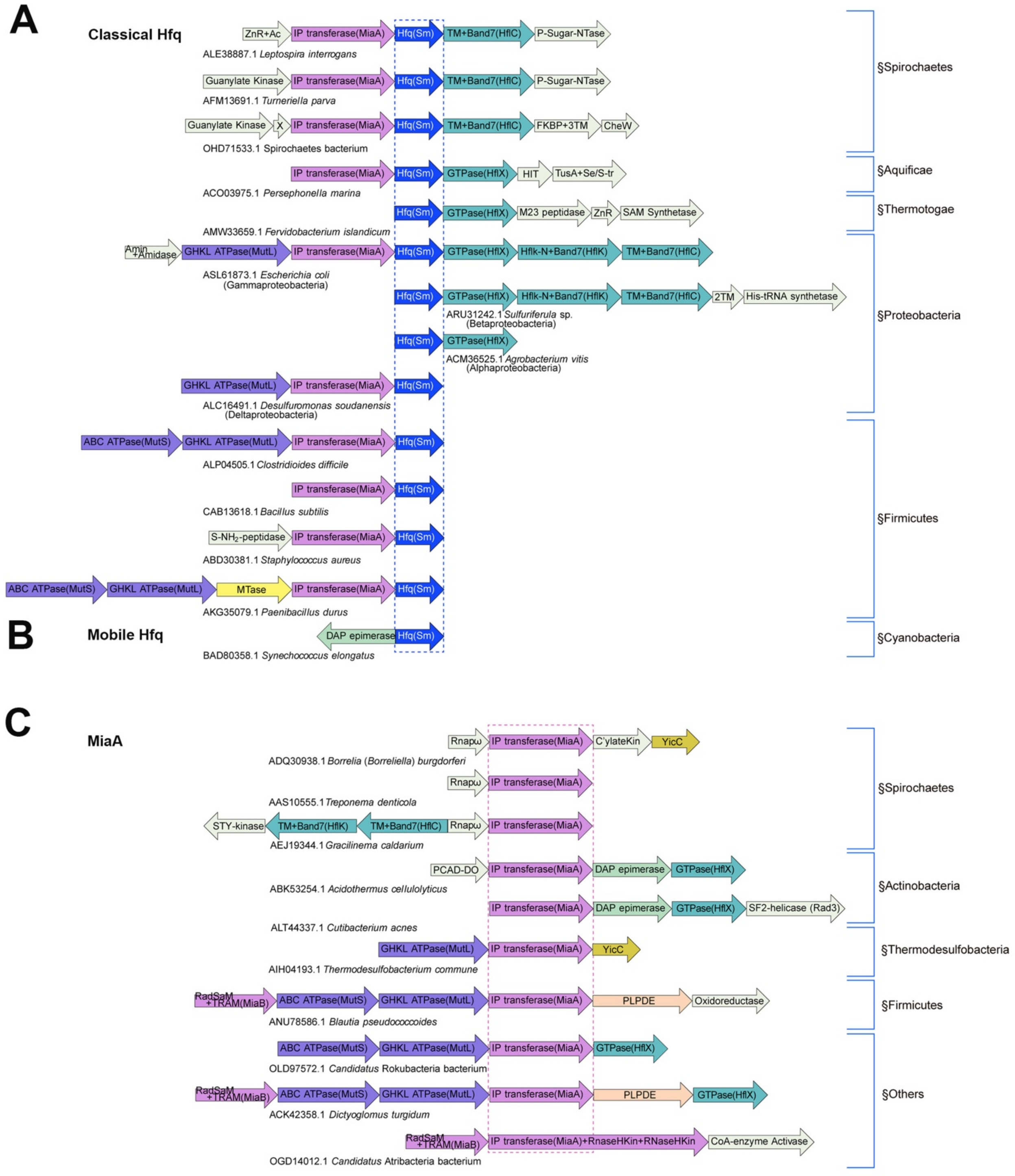


**Supp. Figure S2. The conserved *hfq* and *miaA* gene neighborhoods** (for Figure 1).

Gene neighborhoods from representative species from across the bacterial tree of life for:

(**A**) The classical *hfq*, the homohexameric RNA binding protein of the Sm domain superfamily, clade;

(**B**) The mobile *hfq* clade;

(**C**) The *miaA*, the tRNA adenine isopentenyl transferase, clade.

For panels A–C: the genes are shown as box arrows (only drawn to approximate size) and labeled with the domain architectures of the encoded proteins. Each operon is labeled, below the box arrows, based on the GenBank accession of the anchor gene and the species name. All distinct core genes of the neighborhood are colored differently, and the anchor genes are marked with a dashed box. The genes specific to particular operons but not otherwise widely conserved are colored off-white.

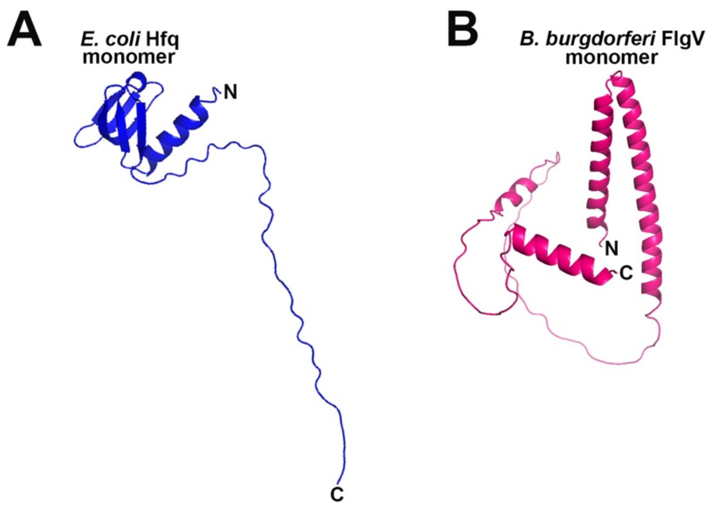


**Supp. Figure S3. AlphaFold2 secondary structure predictions for *E. coli* Hfq and *B. burgdorferi* BB0268 (FlgV)** (for Figure 1). Amino acid sequences analyzed by AlphaFold2 (Jumper *et al.*, 2021) for:

(**A**) *E. coli* Hfq;

(**B**) *B. burgdorferi* BB0268 (FlgV).

For panels A and B: structure figures of the AlphaFold2 output with the highest confidence were prepared using PyMol.

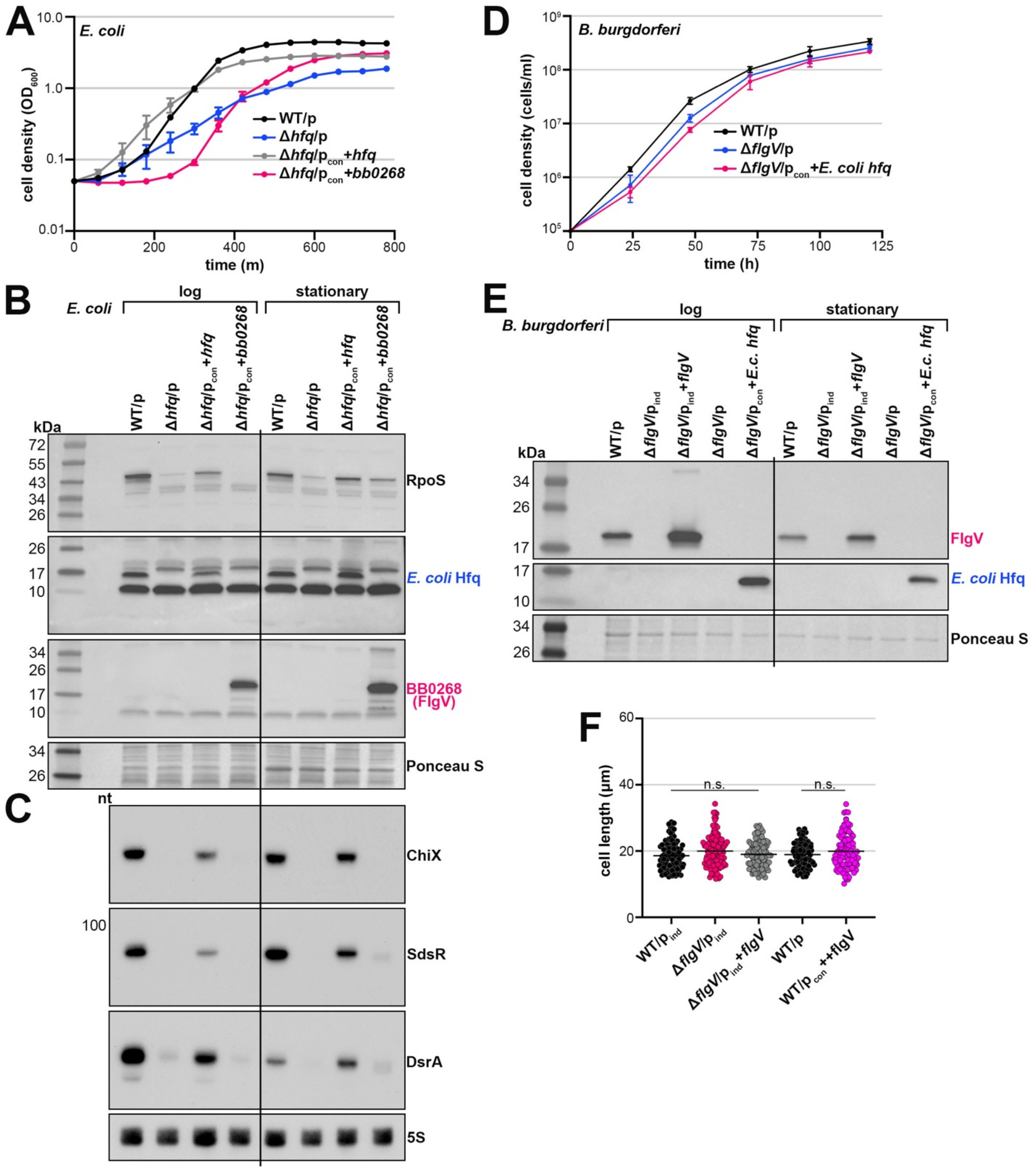


**Supp. Figure S4. BB0268 (FlgV) does not complement an *E. coli* Δ*hfq* strain and *E. coli* Hfq does not complement a *B. burgdorferi* Δ*bb0268* (*flgV*) strain** (for Figures 1 and 3).

(**A**) Growth curve of Δ*hfq* E. coli expressing *bb0268*. WT (EC013) or Δ*hfq* (EC023) *E. coli* were freshly transformed with the empty vector, $p_{con}$+*hfq* and/or $p_{con}$+*bb0268*. Single colonies were isolated and grown overnight. Cell growth was monitored by $OD_{600}$ every 60 min for 780 min total, after a dilution of the overnight culture to an $OD_{600}$ of 0.05. Each data point (circles) represents the mean of three biological replicates with the standard deviation.

(**B**) *E. coli* cultures, defined in panel A, were grown and harvested at log phase (300 min for WT/p and Δ*hfq*/$p_{con}$+*hfq*; 360 min for Δ*hfq*/p and Δ*hfq*/$p_{con}$+*bb0268*) and stationary phase (600 min for WT/p and Δ*hfq*/$p_{con}$+*hfq*; and 720 min for Δ*hfq*/p and Δ*hfq*/$p_{con}$+*bb0268*). Protein extracts were separated on a Tris-Glycine gel, transferred to a membrane, stained with Ponceau S as a loading control, and probed using α-RpoS, α-*E.coli*-Hfq, and α-BB0268(FlgV) antibodies. Proteins were probed sequentially on the same membrane; size markers are indicated.

(**C**) Northern analysis of the *E. coli* ChiX, SdsR and DsrA sRNAs. RNA extracts isolated from the same cultures in panel B were separated on an acrylamide gel, transferred to a membrane and probed for specific sRNAs (RNAs were probed sequentially on the same membrane) and 5S as a loading control. Size markers are indicated for all RNAs, except where the portion of the blot shown is less than 100 nt.

(**D**) Growth curve of Δ*flgV*(*bb0268*) *B. burgdorferi* expressing *E. coli* Hfq. Cell growth was monitored by dark field microscopy enumeration at the indicated time points, after dilution of the starter cultures (PA023, PA257, PA261) to $1x10^5$ cells/ml. Each data point (circles) represents the mean of three biological replicates with the standard deviation.

(**E**) Immunoblot analysis of BB0268 and *E. coli* Hfq levels in *B. burgdorferi*. WT/$p_{ind}$ (PA273), Δ*flgV*/$p_{ind}$ (PA310), and Δ*flgV*/$p_{ind}$+*flgV* (PA312) were grown with 0.1 mM IPTG, after dilution of the starter culture, to an average density of $\sim 1.1x10^7$ cells/ml (log) and $\sim 2.3x10^8$ cells/ml (stationary), and total protein was isolated. Δ*flgV*/p (PA257) and Δ*flgV*/$p_{con}$+*E.coli*(*E.c.*) *hfq* (PA261) were grown, after dilution of the starter culture, to an average density of $\sim 1.4x10^7$ cells/ml (log) and $\sim 2.8x10^8$ cells/ml (stationary), and total protein was isolated. Δ*flgV*/$p_{ind}$ (PA310), Δ*flgV*/p (PA257) and Δ*flgV*/$p_{con}$+*E.coli*(*E.c.*) *hfq* (PA261) samples were collected 9 h after the WT/$p_{ind}$ (PA273) and Δ*flgV*/$p_{ind}$+*flgV* (PA312) samples, so that all cultures were collected at the same cell density. Protein extracts were separated on a Tris-Glycine gel, stained with Ponceau S as a loading control, and probed with α-FlgV and α-*E.coli*-Hfq antibodies. A subset of the lanes are shown in Figure 3A. Proteins were probed sequentially on the same membrane; size markers are indicated.

(**F**) Quantification of spirochete length, for the strains in Figure 3D–I, at stationary phase. All cultures were grown to an average density of ~$1.8 \times 10^8$ cells/ml, washed with 1X PBS and imaged. Approximately 100 cells were traced using the curve (spline) tool with ZEN 3.4 (blue edition) software. Each data point (circles) represents the length of one spirochete; the line indicates the mean length for each strain. Average length across WT/$p_{ind}$, Δ*flgV*/$p_{ind}$, and Δ*flgV*/$p_{ind}$+*flgV* samples or WT/p and WT/$p_{con}$++*flgV* were compared by one-way ANOVA with Tukey's multiple comparisons test or t test, respectively, GraphPad Prism 9.5.1 (n.s., not significant).

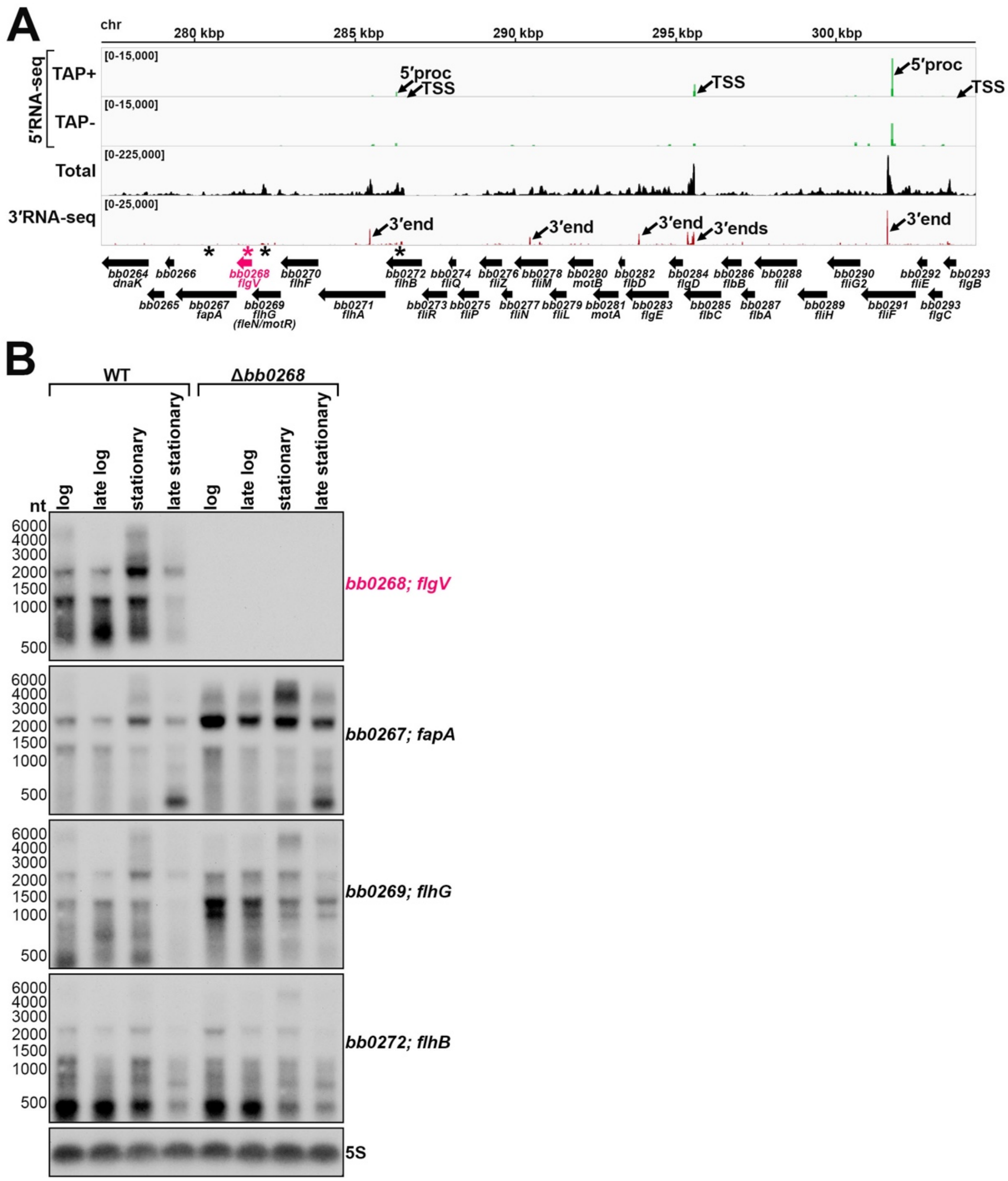


**Supp. Figure S5. *bb0268* (*flgV*) is co-transcribed with flagellar genes** (for Figure 2).

(**A**) RNA-seq browser image of the "flgB superoperon", including the *bb0268* (*flgV*) locus. Browser images display sequencing reads for one DNA strand for RNA isolated from logarithmic phase cells. 5′RNA-seq tracks (green) represent an overlay of two biological replicates (Adams *et al.*, 2017), while total (black) and 3′RNA-seq (red) tracks represent an overlay of three biological replicates (Petroni *et al.*, 2023). Read count ranges are shown in the top left of each frame. The chromosomal coordinates and relative orientation of ORFs (wide black arrows) are depicted. Select transcription start sites (TSS; as determined by the ratio of reads between ±TAP tracks), 5′ processed ends (5′proc) and 3′ ends are indicated (small black arrows).

(**B**) Northern analysis of *bb0268* (*flgV*) across growth. Strains PA001 and PA011 were harvested at time points after the dilution of the subculture for spirochetes in logarithmic phase (log): WT $1.6 \times 10^7$ cells/ml, 36 h; Δ*bb0268* $1.5 \times 10^7$ cells/ml, 48 h; late logarithmic phase (late log): WT $7.5 \times 10^7$ cells/ml, 48 h; Δ*bb0268* $7.8 \times 10^7$ cells/ml, 57 h; stationary phase (stationary): WT $1.7 \times 10^8$ cells/ml, 72 h; Δ*bb0268* $1.9 \times 10^8$ cells/ml, 81 h; late stationary phase (late stationary): WT $3.7 \times 10^8$ cells/ml, 96 h; Δ*bb0268* $1.7 \times 10^8$ cells/ml, 105 h. For each timepoint, total RNA was isolated, separated on an agarose gel, transferred to a membrane and probed for the indicated RNAs. Approximate probe sequence location is indicated by the pink (*bb0268*) or black (*bb0267*; *bb0269*, *bb0272*) asterisks on the corresponding browser image (panel A); size markers are indicated. RNAs were probed sequentially on the same membrane. The membrane was also probed for 5S as a loading control.

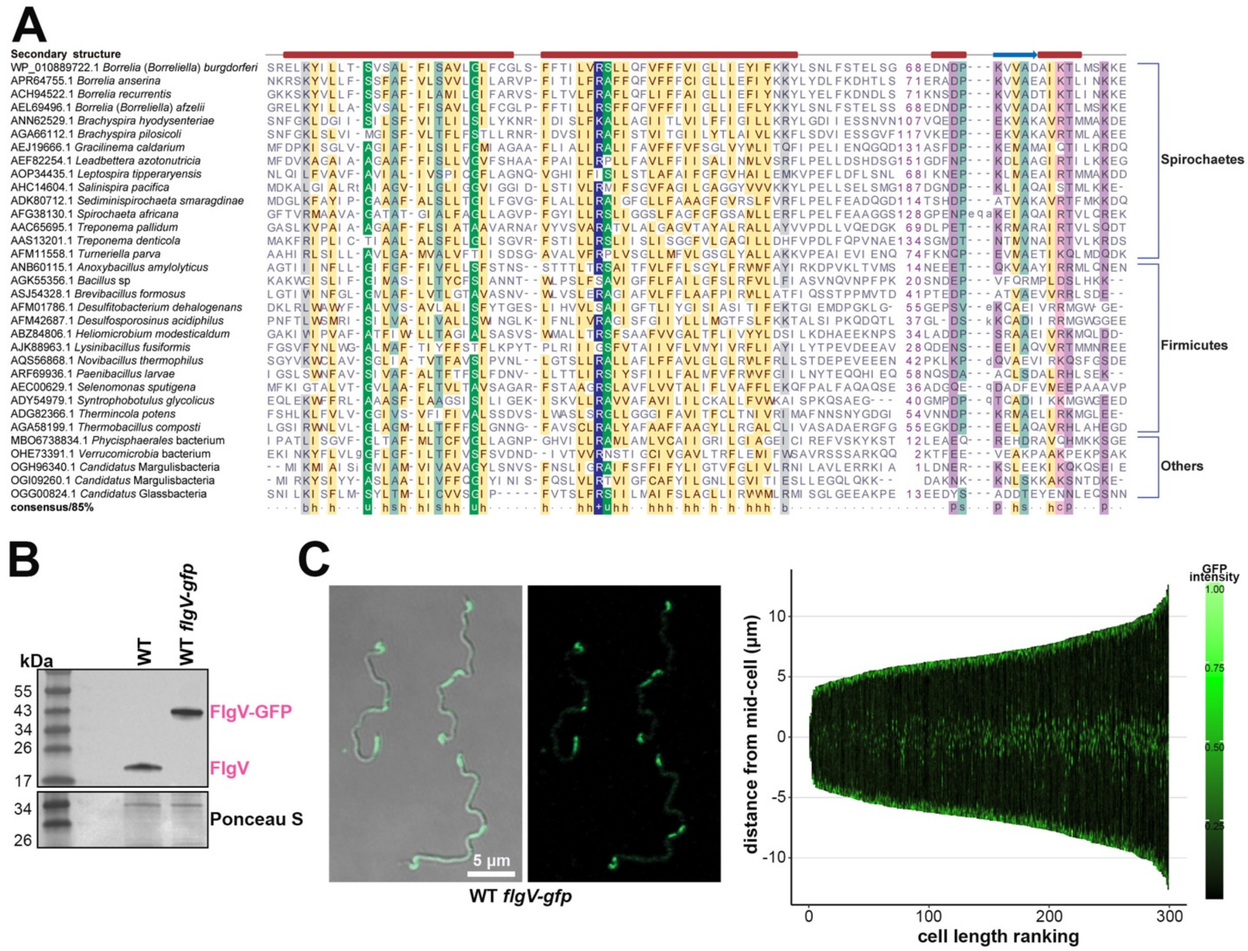


**Supp. Figure S6. FlgV is a conserved, membrane-associated protein** (for Figures 2 and 4)

**(A)** A multiple sequence alignment of the newly identified Type-1 FlgV clade from spirochetes, firmicutes (Bacillota) and other lineages. The sequences are labeled using their GenBank accession and species name and colored as per the 85% consensus using the following amino acid side chain classes in the following order of priority: +: positive, c: charged, u: tiny, l: aliphatic, h: hydrophobic, p: polar, s: small, b: big. The predicted secondary structure is shown above the alignment.

**(B)** Immunoblot analysis of *B. burgdorferi* expressing *gfp*. Total protein was isolated from WT (PA001) and *flgV-gfp* (PA402) *B. burgdorferi*, and protein extracts were separated on a Tris-Glycine gel, stained with Ponceau S as a loading control, and probed with α-FlgV antibodies. Size markers are indicated.

(**C**) Localization of FlgV-GFP by confocal microscopy. The same culture of *B. burgdorferi* (PA402) in Figure 4B was grown to a density of ~$2.6 \times 10^8$ cells/ml, prior to imaging. Left panels: Bright field and fluorescence composite (left micrograph) and fluorescence-only (right micrograph) are shown. Scale bar on left micrograph also applies to right micrograph. Right panel: Demograph of the GFP profile for 300 spirochetes, cells were positioned in order of increasing length.

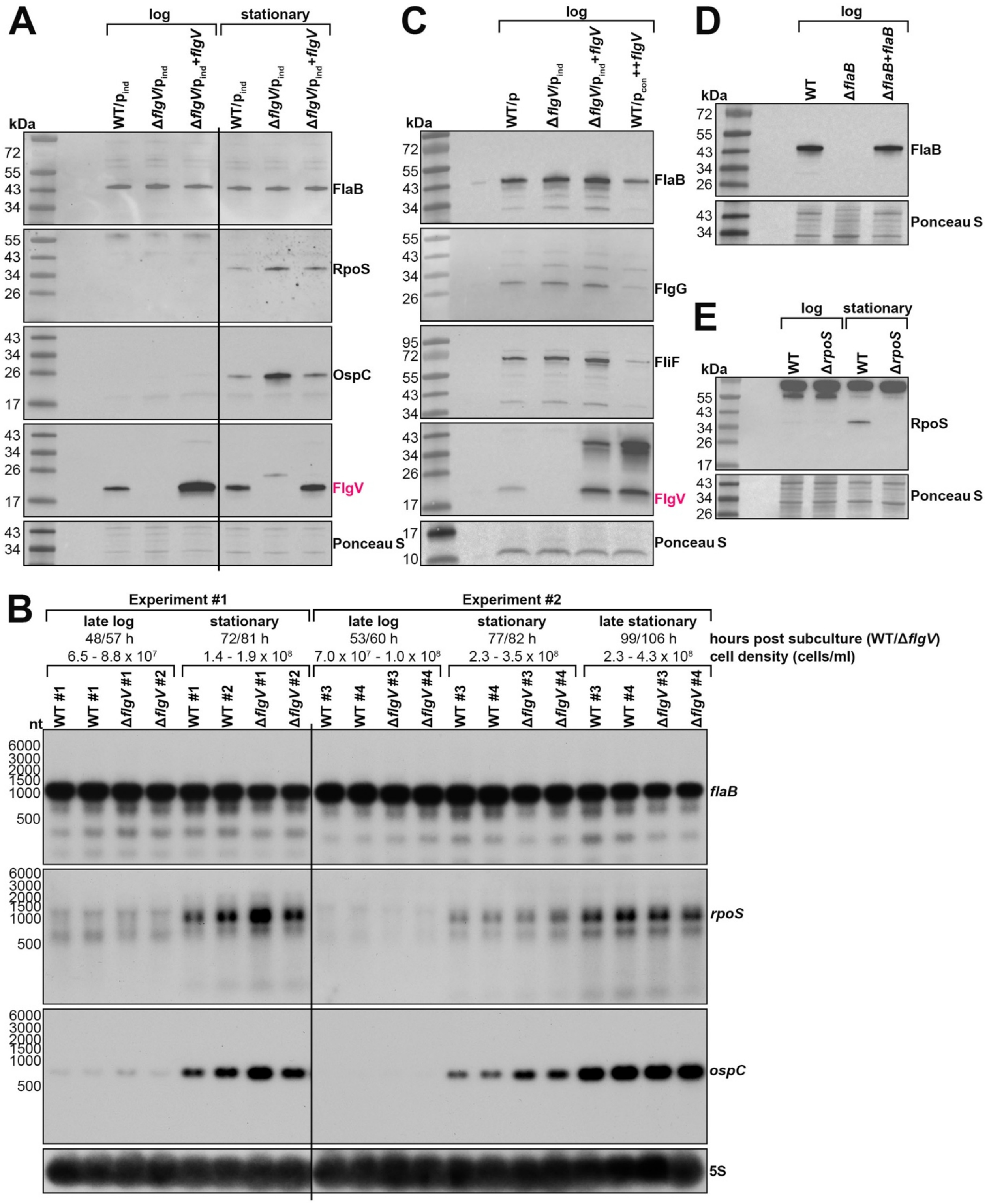


**Supp. Figure S7. FlgV levels impact FlaB protein, but not *flaB* mRNA levels** (for Figure 5).

(**A**) Immunoblot analysis of FlaB and RpoS when FlgV is deleted. Protein extracts from the same samples in Figure 3A were separated on an independent Tris-Glycine gel, stained with

Ponceau S as a loading control, and probed with α-FlaB (Barbour *et al.*, 1986), α-RpoS (this study; see panel E), α-OspC and α-FlgV antibodies. Proteins were probed sequentially on the same membrane; size markers are indicated.

**(B)** Northern analysis of the *flaB*, *rpoS*, and *ospC* mRNAs. RNA extracts isolated from two independent experiments, each with two biological replicates of WT (PA001) and Δ*bb0268* (PA011) cultures, were separated on an agarose gel, transferred to a membrane and probed for specific mRNAs (RNAs were probed sequentially on the same membrane) and 5S as a loading control. The cell density and time of cell collection (WT/Δ*flgV*), after a subculture to $1x10^5$ cells/ml, is indicated above the samples. Size markers are indicated.

(**C**) Immunoblot analysis of flagellar structural proteins when *flgV* is deleted or the protein is overproduced. Cells were grown to a density of: $3.75 \times 10^7$ cells/ml for WT/p (PA023), $2.25 \times 10^7$ cells/ml for Δ*flgV*/$p_{ind}$ (PA310), $4.25 \times 10^7$ cells/ml for Δ*flgV*/$p_{ind}$+*flgV* (PA312), and $0.90 \times 10^7$ cells/ml for WT/$p_{ind}$++*flgV* (PA267) *B. burgdorferi*. Cells were lysed by sonication and protein extracts were separated on a Tris-Glycine gel, stained with Ponceau S as a loading control, and probed with α-FlaB (this study; see panel D), α-FlgG, α-FliF, and α-FlgV antibodies. Proteins were probed sequentially on the same membrane; size markers are indicated.

(**D**) Specificity of the FlaB antibody. Cells were grown to a density of: $4.00 \times 10^7$ cells/ml for WT (PA001), $3.00 \times 10^7$ cells/ml for Δ*flaB* (PA365; (Motaleb *et al.*, 2000)), and $1.65 \times 10^7$ cells/ml for Δ*flaB*+*flgV* (PA367; (Sartakova *et al.*, 2001)) *B. burgdorferi*. Protein extracts were separated as described in panel A and probed with α-FlaB (this study).

(**D**) Specificity of the RpoS antibody. Cells were grown to a density of: $4.50 \times 10^7$ cells/ml for WT (PA001) and $3.50 \times 10^7$ cells/ml for Δ*rpoS* (PA235) *B. burgdorferi* for the logarithmic phase samples and $3.0 \times 10^8$ cells/ml for WT (PA001) and $2.65 \times 10^8$ cells/ml for Δ*rpoS* (PA235) *B. burgdorferi* for the stationary phase (48 h later) samples. Protein extracts were separated as described in panel A and probed with α-RpoS (this study).
